# Benchmarking Static Gene Regulatory Network Reconstruction and Dynamic Transition Probing in Single-Cell Foundation Models

**DOI:** 10.64898/2026.05.17.725083

**Authors:** Zhongni Ye, Ning Yang, Xiaojing Yang, Xiaowei Mao, Chao Tang

## Abstract

Single-cell foundation models may encode gene regulatory information, but it remains unclear which model components capture this signal and how it compares with conventional inference methods. Here, we introduce a unified benchmark that evaluates gene regulatory network (GRN) reconstruction from six single-cell foundation models and three classical baselines across six datasets and four reference network types. We disentangle three sources of regulatory signal within each model—pretrained token embeddings, final-layer hidden states, and attention-derived scores. Under a strict zero-shot setting, scGPT token-embedding similarity outperforms classical baselines on STRING and ChIP-seq references, recovers core transcription factors, and best preserves reference network topology. Moreover, static GRNs cannot test whether learned gene–gene relationships are predictive of expression dynamics, we therefore introduce dynamic transition probing, which iteratively applies a model’s reconstruction head to drive early-cell profiles toward late-cell states without temporal supervision. We find pretrained models capture meaningful developmental transitions, with scFoundation showing the strongest overall performance. Together, our results show that single-cell foundation models encode transferable regulatory and dynamical priors, but how well these priors can be recovered depends on model architecture, pretraining design, and extraction strategy.

## 1 Introduction

Gene regulatory networks (GRNs) describe how transcription factors regulate target genes and provide a useful framework for studying cell fate, development, and disease [1, 2]. With the rapid growth of bulk and single-cell transcriptomic data, GRNs can now be inferred across diverse cell types, tissues, and biological states [3, 4]. However, GRN reconstruction remains challenging because transcriptomic data are noisy, high-dimensional, sparse, and strongly context dependent, especially at single-cell resolution [5, 6].

A wide range of computational methods has been developed for GRN reconstruction. Correlation and co-expression methods are simple and scalable, but they cannot distinguish direct regulation from indirect association and are sensitive to batch effects and sparsity [7]. Information-theoretic methods, such as ARACNE [8] and CLR [9], can capture nonlinear dependencies, but their performance depends on sample size and density estimation. Regression-based methods, such as Inferelator [5] and TIGRESS [10], infer regulators for each target gene through sparse or ensemble regression, but they can be affected by noise, collinearity, and linear modeling assumptions. Tree-based and neural methods, such as GENIE3 [11], GRNBoost [12], DeepSEM [13], and DeepDRIM [14], improve model flexibility, but their generalization and interpretability remain difficult to assess. Overall, limited ground truth and strong dataset dependence still make fair evaluation difficult.

Single-cell foundation models provide a new source of information for GRN reconstruction. Models such as Geneformer [15] and scGPT [16] are pretrained on large-scale single-cell datasets and learn representations of both genes and cells, which contain regulatory information that is not easily recovered from a single expression matrix. Instead of directly inferring edges from expression values, one can extract gene representations from pretrained models and construct networks from representation similarity, or use attention scores as model-internal gene–gene interaction signals. These approaches are promising because pretrained models may encode transferable biological priors, reduce noise, and capture context-dependent relationships. However, it remains unclear which model components contain useful GRN information and when these representations actually improve network reconstruction.

A major obstacle is the lack of a unified benchmark for foundation-model-based GRN reconstruction. Standard benchmarks such as DREAM5 [17] and BEELINE [7] have been important for evaluating conventional GRN inference methods, but they were not designed to evaluate the full workflow from foundation-model representations to inferred regulatory networks. Recent zero-shot studies also suggest that pretrained single-cell models do not always transfer reliably across settings [18]. These issues call for a controlled benchmark that compares representation sources, graph-construction strategies, reference networks, and evaluation metrics under the same framework.

In this work, we introduce a unified benchmark for static GRN reconstruction and dynamic transition probing in single-cell foundation models. Our main contributions are:

- **A unified benchmark for foundation-model-based GRN reconstruction**. We evaluate six single-cell foundation models and three classical machine learning baselines across six datasets and four reference networks, using the same preprocessing, graph-construction, and evaluation protocols.
- **A systematic comparison of three representation sources**. We compare three ways of extracting regulatory signals from foundation models: pretrained gene token embeddings, final-layer hidden states, and attention-derived scores.
- **Dynamic transition probing beyond static GRNs**. We introduce a new task that tests whether pretrained generative priors can drive early-cell expression states toward later ones without temporal supervision.

## 2 Benchmark Design and Experimental Setup

### 2.1 Unified benchmark workflow

We build a unified benchmark to test whether representations from single-cell foundation models can support GRN reconstruction. As shown in Figure 1(a), the benchmark follows a fixed pipeline: single-cell expression matrix, representation extraction from pretrained models, GRN construction, and quantitative evaluation. Given the same expression data, all methods are evaluated under the same preprocessing procedure, candidate gene set, graph-construction rules, reference networks, and evaluation metrics.

**Figure 1:**
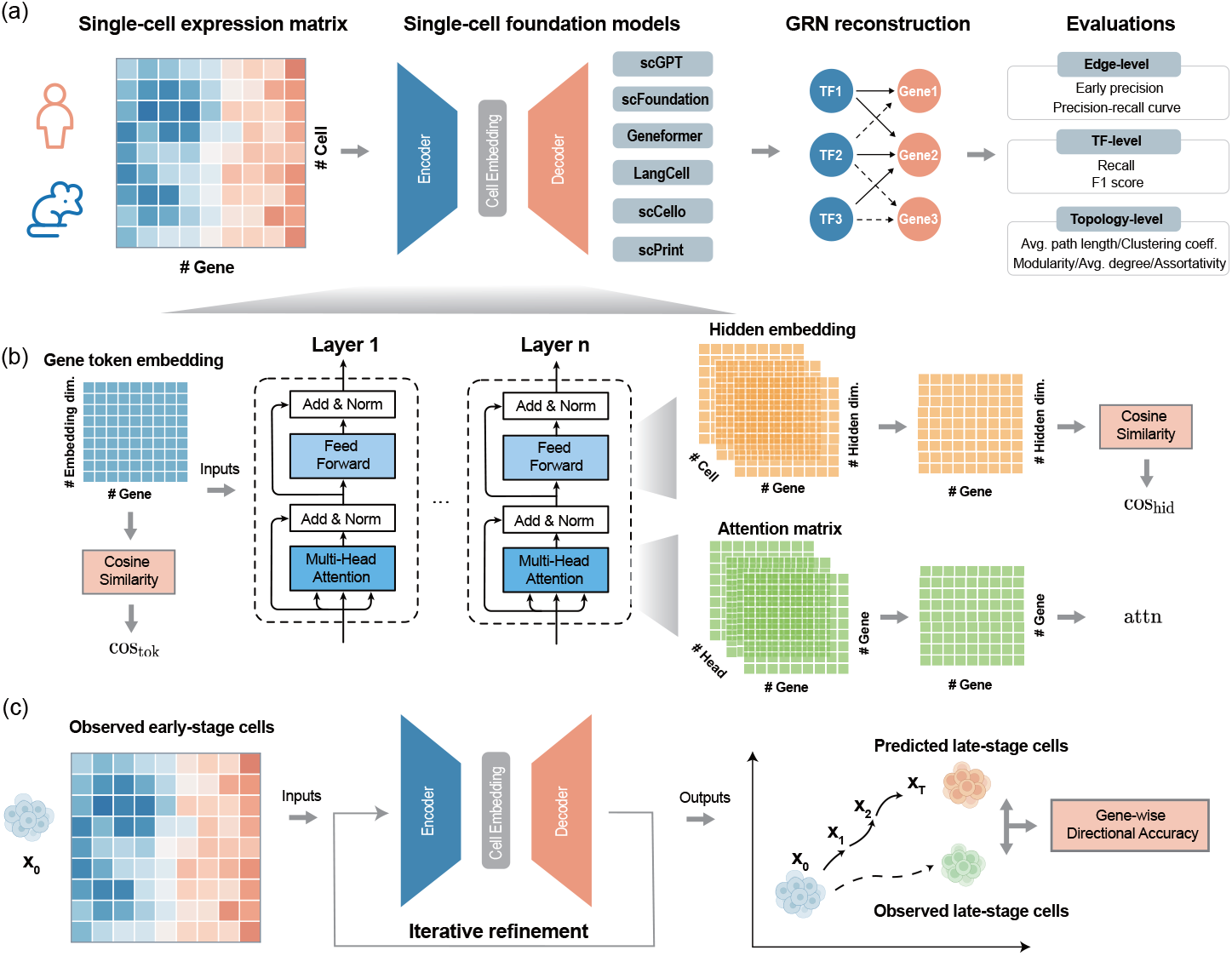
Benchmark workflow for GRN reconstruction and dynamic transition probing. (a) Single-cell expression profiles are processed by pretrained foundation models, and model-derived representations are converted into GRNs for edge-level, TF-level and topology-level evaluation. (b) Three static GRN reconstruction strategies: token-embedding cosine similarity (cos_tok_), hidden-state cosine similarity (cos_hid_), and attention-derived scores (attn). (c) Dynamic transition probing starts from early-stage cell expression profiles and iteratively updates them with a pretrained generative model. The predicted states are compared with observed late-stage cells using gene-wise directional accuracy.

The benchmark uses single-cell expression matrices from six datasets in the BEELINE framework [7]: mouse dendritic cells (mDC) [19], three mouse hematopoietic stem-cell lineages [20]: erythroid (mHSC-E), granulocyte–macrophage (mHSC-GM), and lymphoid (mHSC-L), human mature hepatocytes (hHep) [21], and human embryonic stem cells (hESC) [22]. Raw expression data were preprocessed following the BEELINE protocol. We then constructed two TF-prioritized candidate gene sets: TFs+500 and TFs+1000, which includes all significant TFs plus the top 500/1000 most dynamically variable non-TF genes. Additional dataset details are provided in Appendix A.1.

We evaluate six representative single-cell foundation models, including scGPT [16], scFoundation [23], Geneformer [15], LangCell [24], scCello [25], and scPRINT [26]. These models cover diverse transformer-based architectures, vocabulary designs, embedding dimensions, and pretraining scales. To assess whether foundation-model-derived representations provide advantages over conventional inference methods, we also include three classical machine learning baselines, GENIE3 [11], DeepSEM [13], and GRNboost [27], which infer regulatory relationships directly from expression matrices without pretrained representations. Model details and baseline implementations are provided in Appendix A.2.

The inferred networks are evaluated against four types of reference networks with different evidence sources: cell-type-specific ChIP-seq networks [28, 29, 30], non-specific ChIP-seq networks [31, 32, 33], functional interaction networks from STRING [34], and pathway- or signaling-informed interactions from OmniPath [35]. These references allow us to evaluate whether inferred GRNs recover transcriptional regulation, broader functional association, and signaling-related gene interactions.

We evaluate reconstructed GRNs at three levels. At the edge level, we treat GRN inference as a ranked edge-prediction task and compare predicted gene–gene scores with reference edges using early precision and precision–recall curves. At the TF level, we evaluate whether each method correctly recovers the target genes of individual transcription factors. For each TF, we compare its predicted target set with its reference target set and compute recall and F1 score. At the topology level, we assess whether reconstructed networks preserve global network organization using graph metrics such as average shortest path length, clustering coefficient, modularity, average degree, and assortativity.

### 2.2 Static GRN reconstruction

We compare three representation-to-network strategies for static GRN reconstruction (Figure 1b)). These strategies probe three possible sources of GRN information in single-cell foundation models: static gene-level priors in model parameters, input-dependent contextual representations, and gene– gene coupling implied by the Transformer’s attention mechanism.

- **Pretrained token embeddings (**cos_**tok**_). For each model, we extract gene embeddings from the embedding layer or vocabulary-aligned model parameters. Pairwise gene–gene association scores are then computed via cosine similarity between token embeddings, yielding an undirected GRN. Since these embeddings are independent of any input expression profile, this strategy primarily probes the static biological priors acquired during pretraining.
- **Final-layer hidden states (**cos_**hid**_). We pass each input cell through the model and extract the last-layer hidden representation of each gene token. These cell-wise hidden states are aggregated into gene-level representations, and pairwise cosine similarity is used to construct an undirected GRN. Unlike token embeddings, hidden states are conditioned on the input expression profile and therefore capture context-dependent gene associations.
- **Attention-derived scores (attn)**. For transformer-based models, we extract encoder attention scores of the last layer and aggregate them across heads and cells to obtain gene–gene attention scores. Because attention is asymmetric, the score from gene *i* to gene *j* need not equal the score from gene *j* to gene *i*, naturally producing a directed GRN.

All three strategies output ranked gene–gene scores. For evaluation, we use a unified TF–target candidate space, where the source gene must be a ground-truth TF and the target gene must belong to the reference gene set. For undirected cosine-based networks, the gene–gene similarity score is assigned to the corresponding TF–target pair; for directed attention-derived networks, the original source–target direction is retained. Edges are ranked by absolute weight, and the top-ranked predictions up to the number of ground-truth edges are evaluated using early precision and precision–recall curves. Details are provided in Appendix B.2.

### 2.3 Dynamic transition probing

While static GRN reconstruction tests whether a model encodes gene–gene relationships, a GRN is still an abstract summary of regulatory structure and cannot fully describe how cell states change over time [36]. A more demanding question is whether the learned relationships are functional enough to predict expression dynamics [37, 38], therefore, we introduce dynamic transition probing as a complementary task that directly tests this capacity (Figure 1(c)). This probe is not intended as a supervised forecasting task, instead, it asks whether a pretrained model, when given only an early-cell state, can use its internalized gene–gene dependencies to produce changes that are directionally consistent with later cellular states.

Starting from an observed early-cell state **x**^(0)^, each model is iteratively updated to generate a sequence of predicted states, which we treat as a model-induced transition path **x**^(0)^, **x**^(1)^, …, **x**^(*T*)^. The update rule depends on the model input format: for continuous-value models, such as scGPT, scFoundation, and scPRINT, the current expression state is passed through the pretrained model to predict a refined expression profile; for rank-driven models, such as Geneformer, LangCell, and scCello, we refine the gene-rank ordering through likelihood-improving token swaps while preserving the same expressed gene set. In both cases, repeated updates produce a terminal predicted state starting from the early-cell state.

We evaluate whether the predicted changes follow the observed early-to-late direction. Specifically, we compare the sign of the predicted expression or rank change with the sign of the observed expression change, and report directional accuracy on the top 30% genes with the largest true change magnitude. Full implementation details are provided in Appendix C.

## 3 Results

### 3.1 scGPT token embeddings enable strong zero-shot GRN reconstruction

We first evaluated static GRN reconstruction on the TFs+500 benchmark. We compared six single-cell foundation models and three classical machine learning baselines across six single-cell datasets. Performance was measured by the early precision ratio (EPR) and the area under the precision–recall curve ratio (AUPRC ratio). For each method, results were averaged across the six datasets and three main reference sources: STRING, non-specific ChIP-seq, and OmniPath. Cell-type-specific ChIP-seq references were not included in the main comparison because all methods showed uniformly near-random performance on this setting. We also repeated the analysis on the larger TFs+1000 benchmark and observed similar conclusions; complete results are provided in Appendix E.

As shown in Table 1, scGPT with token-embedding cosine similarity (cos_tok_) achieved the strongest overall performance among foundation-model-based methods. It obtained the best EPR and AUPRC ratio on STRING and non-specific ChIP-seq references, and also performed well on OmniPath. Although the standard deviations are sizable because performance varies across datasets, scGPT cos_tok_ consistently ranked first or near-first on individual datasets, as shown in the per-dataset results in Table 4& 6.

**Table 1:** Performance on the TFs+500 benchmark across six single-cell datasets and three reference networks (STRING, Cell-type-specific ChIP-seq, OmniPath). Values are mean ± standard deviation across datasets. Both metrics are reported as ratios over a random predictor; values *>*1 indicate better-than-random performance. Bold marks the best value in each column.

| Model | Method | Early precision ratio |  |  | AUPRC ratio |  |  |
| --- | --- | --- | --- | --- | --- | --- | --- |
|  |  | STRING | ChIP-seq | OmniPath | STRING | ChIP-seq | OmniPath |
| **scGPT** | cos<sub>tok</sub> | **6.568±1.772** | **4.211±1.703** | 2.387±0.603 | **5.632±2.310** | **3.002±1.294** | 1.460±0.242 |
| | attn | 1.255±0.548 | 1.299±0.290 | 0.998±0.334 | 0.941±0.064 | 0.990±0.076 | 1.000±0.044 |
| | cos<sub>hid</sub> | 4.301±0.983 | 2.266±0.829 | 1.425±0.667 | 3.104±1.130 | 1.563±0.480 | 1.141±0.158 |
| **scFoundation** | cos<sub>tok</sub> | 1.174±0.051 | 1.223±0.309 | 1.150±0.106 | 1.211±0.147 | 1.144±0.129 | 1.018±0.030 |
| | attn | 0.956±0.196 | 0.997±0.334 | 1.091±0.245 | 1.017±0.062 | 1.016±0.041 | 1.025±0.042 |
| | cos<sub>hid</sub> | 1.127±0.392 | 1.025±0.392 | 0.997±0.811 | 1.097±0.247 | 1.033±0.076 | 1.010±0.051 |
| **Geneformer** | cos<sub>tok</sub> | 1.821±0.546 | 0.932±0.555 | 1.009±0.721 | 1.244±0.157 | 1.077±0.178 | 1.015±0.266 |
| | attn | 3.082±1.157 | 1.644±0.771 | 1.203±0.513 | 1.867±0.635 | 1.175±0.239 | 1.083±0.085 |
| | cos<sub>hid</sub> | 1.523±0.457 | 0.983±0.381 | 0.788±0.420 | 1.304±0.191 | 1.065±0.127 | 0.969±0.130 |
| **LangCell** | cos<sub>tok</sub> | 1.837±0.498 | 0.970±0.570 | 1.002±0.711 | 1.251±0.163 | 1.080±0.182 | 1.016±0.263 |
| | attn | 2.104±0.910 | 1.343±0.571 | 1.547±1.072 | 1.591±0.330 | 1.129±0.201 | 1.176±0.384 |
| | cos<sub>hid</sub> | 2.133±0.455 | 1.694±0.389 | 1.523±0.343 | 1.584±0.132 | 1.276±0.166 | 1.203±0.101 |
| **scCello** | cos<sub>tok</sub> | 2.194±0.573 | 1.477±0.498 | 1.450±0.177 | 1.502±0.430 | 1.120±0.100 | 1.117±0.036 |
| | attn | 1.289±0.168 | 1.307±0.207 | 2.132±1.124 | 1.262±0.173 | 1.092±0.143 | 1.345±0.300 |
| | cos<sub>hid</sub> | 2.578±0.638 | 1.936±0.790 | 1.896±0.434 | 1.676±0.265 | 1.204±0.191 | 1.226±0.182 |
| **scPRINT** | cos<sub>tok</sub> | 0.948±0.228 | 0.798±0.265 | 0.662±0.171 | 0.980±0.042 | 0.976±0.095 | 0.887±0.057 |
| | attn | 0.920±0.472 | 1.074±0.400 | 1.071±0.380 | 1.217±0.173 | 1.127±0.126 | 1.085±0.154 |
| | cos<sub>hid</sub> | 1.381±0.446 | 1.291±0.441 | 1.402±0.409 | 1.281±0.201 | 1.155±0.071 | 1.115±0.093 |
| **ML Baselines** | GENIE3 | 4.656±2.500 | 3.025±1.670 | 1.575±0.288 | 3.301±2.244 | 1.793±0.848 | 1.175±0.181 |
| | DeepSEM | 5.391±2.635 | 3.789±1.936 | 2.167±0.433 | 4.087±2.622 | 2.338±1.196 | 1.305±0.137 |
| | GRNBoost | 4.121±1.949 | 2.680±1.445 | **3.781±1.365** | 1.257±0.172 | 1.462±0.259 | **2.262±0.729** |

Classical machine learning baselines remained competitive, but they fit a dataset-specific model on each test expression matrix, while cos_tok_ uses only fixed pretrained token embeddings without any dataset-specific updates. Even under this stricter zero-shot setting, scGPT cos_tok_ outperformed all three classical baselines on STRING and non-specific ChIP-seq, indicating that the pretrained embedding space already encodes a transferable regulatory signal. The exception is OmniPath, where GRNBoost remained the best, suggesting that dataset-specific fitting is still advantageous for some reference networks.

The three extraction strategies behaved differently across models. For scGPT, cos_tok_ clearly outperformed cos_hid_ and attn, suggesting that regulatory information is strongly encoded in its token embedding space. Geneformer performed better with attention-based scores, suggesting that self-attention may be a more informative source for GRN reconstruction in this model. In contrast, scCello and LangCell often benefited more from hidden-state representations, suggesting that their additional cell-level or cross-modal training objectives may place more regulatory signal in contextual embeddings. These results suggest that the best extraction strategy depends on the model architecture and pretraining design.

### 3.2 Topological fidelity of reconstructed GRNs

We next asked whether reconstructed GRNs preserve the global topology of reference regulatory networks. This analysis goes beyond edge-level accuracy by testing whether a method recovers broader network organization. We compared each reconstructed network with the STRING reference using five classical topological metrics [39]: average shortest path length, clustering coefficient, modularity, average degree, and assortativity. Implementation details are provided in Appendix D.3. Figure 2 shows hESC as a representative dataset, with results on other datasets reported in Table 8. As shown in Figure 2(a)–(c), embedding-based methods generally produced networks whose topology was closer to the STRING reference than attention-based methods. This was observed for both token-embedding similarity (cos_tok_) and hidden-state similarity (cos_hid_), suggesting that embedding spaces better preserve the global organization of GRNs. In contrast, attention-derived networks showed larger deviations across multiple topological metrics.

**Figure 2:**
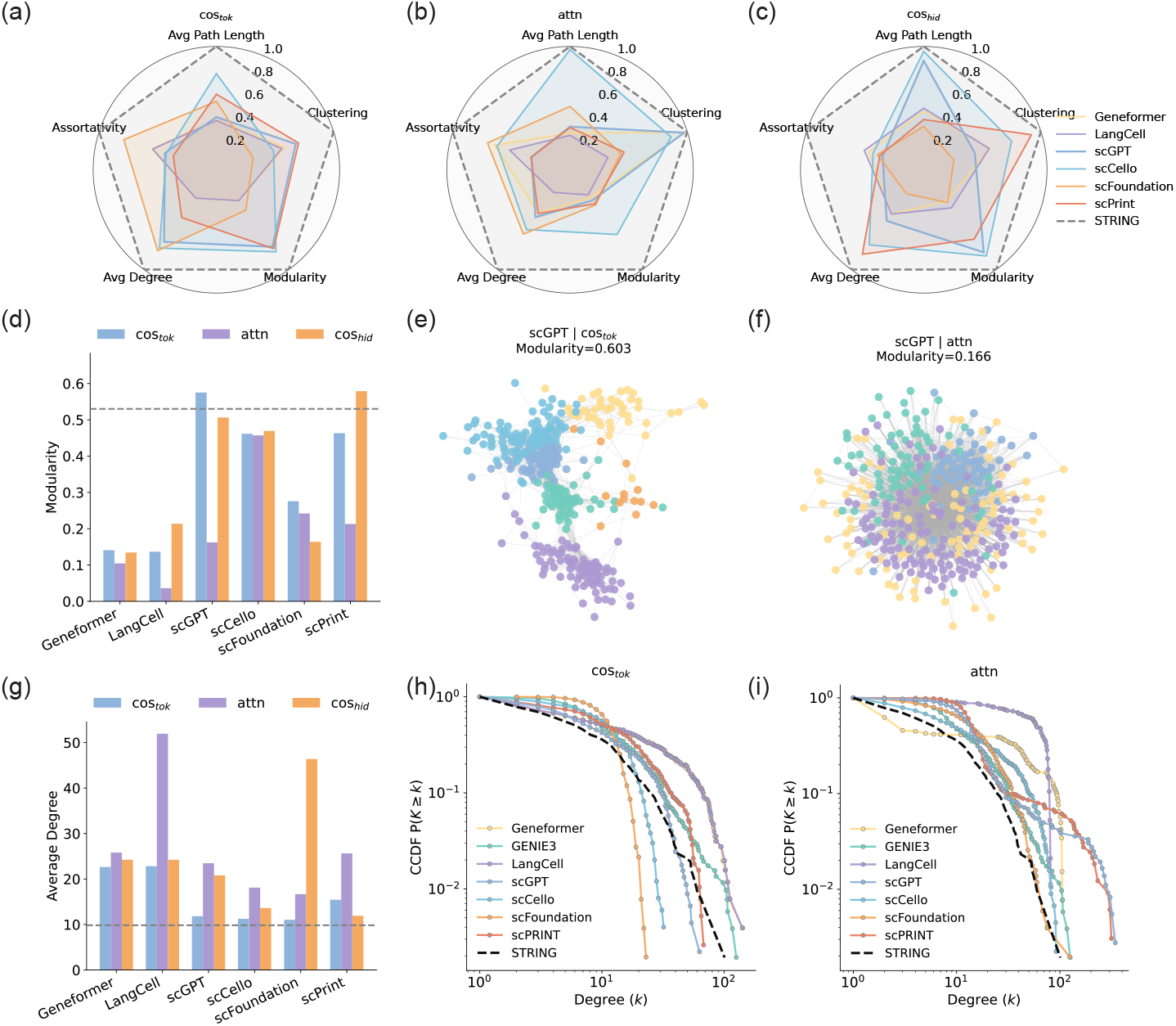
Topological analysis of reconstructed GRNs on the hESC dataset. (a)–(c) Comparison of reconstructed networks across five topological metrics for cos_tok_, attn, and cos_hid_. The dashed line represents the STRING reference network, and values closer to the dashed line indicate higher topological fidelity. (d) Modularity comparison of the three extraction strategies across six foundation models. (e)–(f) Network visualization of scGPT GRNs reconstructed using cos_tok_ and attention scores. (g) Average Degree comparison of the three extraction strategies across six foundation models. (h-i) Degree CCDF curves of networks reconstructed by cos_tok_ and attn, compared with the STRING reference.

The difference was especially clear in modularity. Attention-based networks often had lower modularity than embedding-based networks, indicating weaker community structure (Figure 2(d)). In scGPT, this was also visible in the network layouts: the cos_tok_ network showed clearer and more separated communities, whereas the attention-based network appeared more mixed and less organized (Figure 2(e),(f)). To further examine this limitation, we constructed separate networks from each scGPT attention head and computed their topological metrics (Figure 6). These head-specific networks varied substantially in modularity, suggesting that only some heads preserve useful community structure. Thus, directly aggregating all attention heads may dilute topological signals relevant to GRN reconstruction.

Finally, we examined whether reconstructed networks preserve the degree structure of STRING. Many reconstructed networks, especially attention- or hidden-state-based networks, had higher average degree than the STRING reference, suggesting that they tend to produce overly dense graphs (Figure 2(g)). We further compared degree distributions using the complementary cumulative distribution function (CCDF), which captures heavy-tailed structure with many low-degree genes and a few highly connected hubs. As shown in Figure 2(h)–(i), cos_tok_ networks generally followed the STRING degree-decay pattern more closely, especially scGPT cos_tok_, whereas attention-based networks showed larger deviations. These results suggest that token-embedding similarity better preserves the hub-dominated degree structure of reference GRNs.

### 3.3 scGPT token embeddings recover high-confidence edges and core TFs

Since scGPT cos_tok_ showed the strongest overall performance, we further compared the three scGPT extraction strategies in detail. We first asked whether true regulatory edges are enriched among top-ranked predictions. As shown in Figure 3(a), cos_tok_ recovered a much higher fraction of true edges than cos_hid_ and attn under high-confidence thresholds. This indicates that true regulatory interactions are more concentrated at the top of the cos_tok_ ranking, which is important in practice because experimental validation usually focuses on a small number of high-confidence candidates.

**Figure 3:**
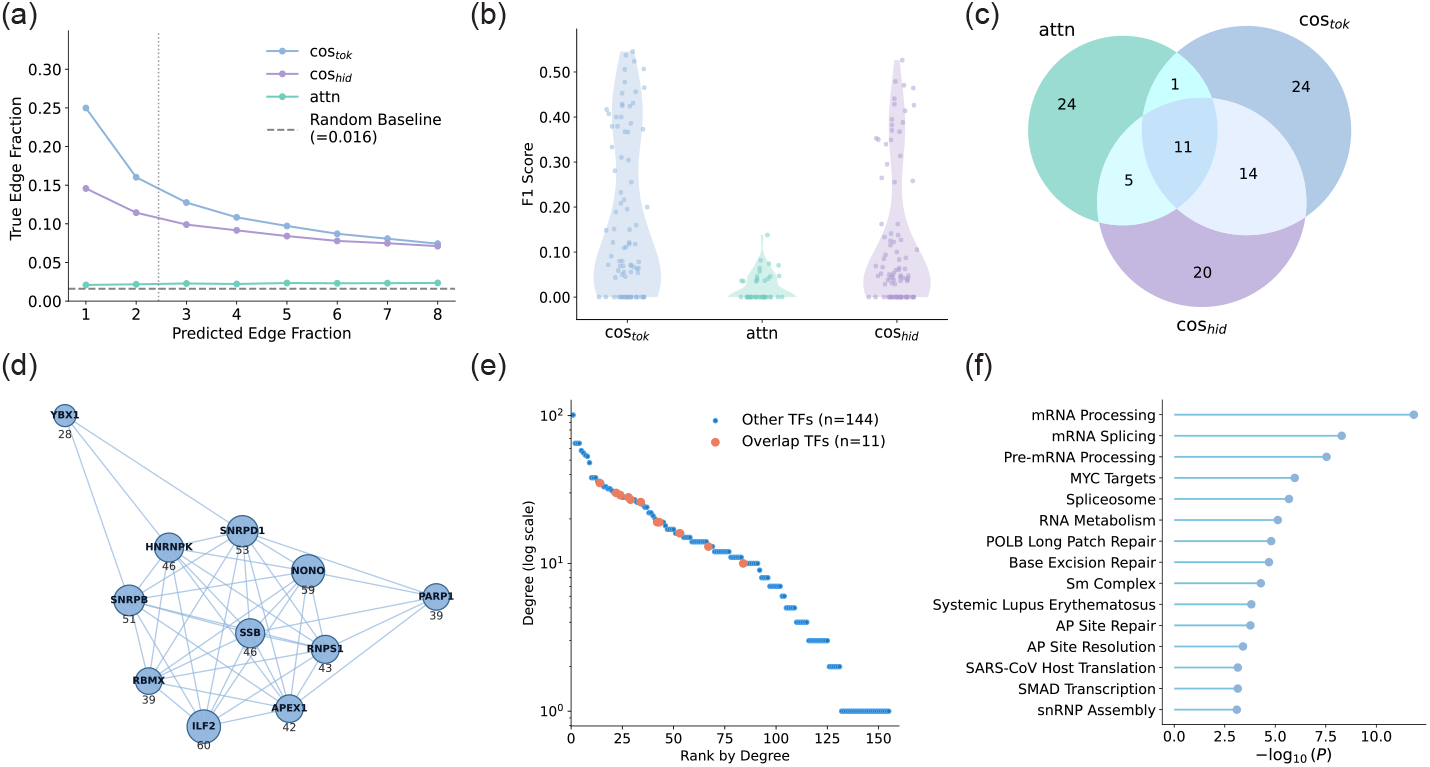
Detailed comparison of scGPT extraction strategies. (a) Fraction of true regulatory edges among top-ranked predictions under different prediction thresholds. Horizontal and vertical dashed line marks the random baseline and the threshold used for early precision. (b) Distribution of TF-level F1 scores for core TFs identified by cos_tok_, cos_hid_, and attn. (c) Overlap of core TF sets identified by the three extraction strategies on the hESC dataset. (d) Interaction network of consensus TFs shared by all three strategies. Node size is proportional to degree, with degree values shown below nodes. (e) Degree-rank plot of the union of predicted core TFs, with the consensus TFs highlighted. (f) Functional enrichment of consensus TFs. Terms are ranked by enrichment significance, measured as − log_10_(*P*); larger values indicate stronger enrichment.

We next evaluated whether each strategy can identify important TFs. This tests GRN quality at the regulator level: beyond recovering correct edges, a useful network should also highlight key regulators. TF-level F1 scores were computed from predicted regulator sets, with details provided in Appendix B.3. As shown in Figure 3(b), cos_tok_ and cos_hid_ identified TFs with substantially higher F1 scores, although their distributions were heterogeneous. In contrast, attn scores were mostly concentrated near zero, suggesting that direct attention scores provide weaker TF-level signal in scGPT.

Finally, we compared the regulators selected by different strategies on the hESC dataset. As shown in Figure 3(c), cos_tok_ and cos_hid_ showed the largest pairwise overlap, whereas attn identified many strategy-specific TFs. We then focused on the 11 consensus TFs shared by all three strategies. These TFs formed a connected interaction network and ranked near the top by degree, suggesting that they are consistently recovered core regulators (Figure 3(d),(e)). Functional enrichment analysis showed that these consensus TFs were enriched in RNA-related processes, including mRNA processing, RNA splicing, spliceosome-associated pathways, and DNA repair (Figure 3(f)). Together, these results suggest that overlap across extraction strategies highlights a robust core of biologically meaningful regulators, while strategy-specific TFs may reflect representation-dependent signals.

### 3.4 Dynamic transition probing reveals directional expression dynamics in pretrained models

Beyond static GRN reconstruction, we asked whether single-cell foundation models have internalized gene–gene dependencies that can support dynamic expression changes without temporal supervision. Using the iterative refinement framework described in Sec. 2.3, we initialized each model with early-cell states and examined whether the induced updates moved them toward later cell states across six datasets. Performance was measured by directional accuracy on the top 30% genes with the largest observed early-to-late changes, with full details provided in Appendix C.

We first used scGPT on the mHSC-L dataset as a representative example. As shown in Figure 4(a), the predicted terminal states shifted from the early-cell population toward the observed late-cell population, suggesting that the model-induced updates capture a meaningful developmental direction rather than random fluctuations. During iterative refinement, directional accuracy increased over successive steps and then gradually plateaued for most models (Figure 4(b)), indicating that repeated model updates progressively improve the predicted transition states.

**Figure 4:**
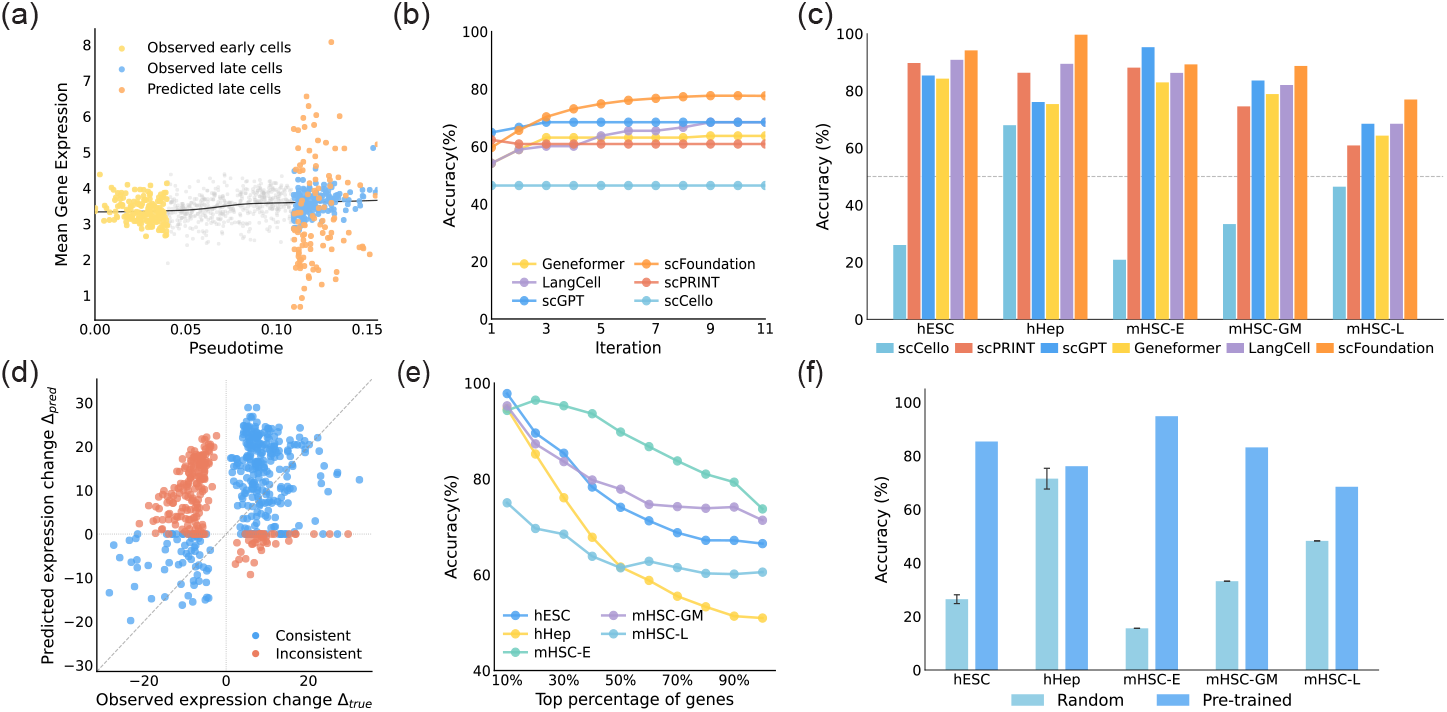
Dynamic transition probing with single-cell foundation models. (a) Representative scGPT prediction on the mHSC-L dataset, showing observed early cells, observed late cells, and predicted terminal states in gene expression level. (b) Directional prediction accuracy during iterative refinement for different models on mHSC-L. (c) Directional prediction accuracy across five datasets and six models. (d) Relationship between observed expression changes and predicted expression changes for scGPT on mHSC-L. (e) Directional prediction accuracy as a function of gene expression-change magnitude for scGPT on mHSC-L. (f) Comparison between pretrained and randomly initialized scGPT weights under the same iterative refinement procedure.

**Figure 5:**
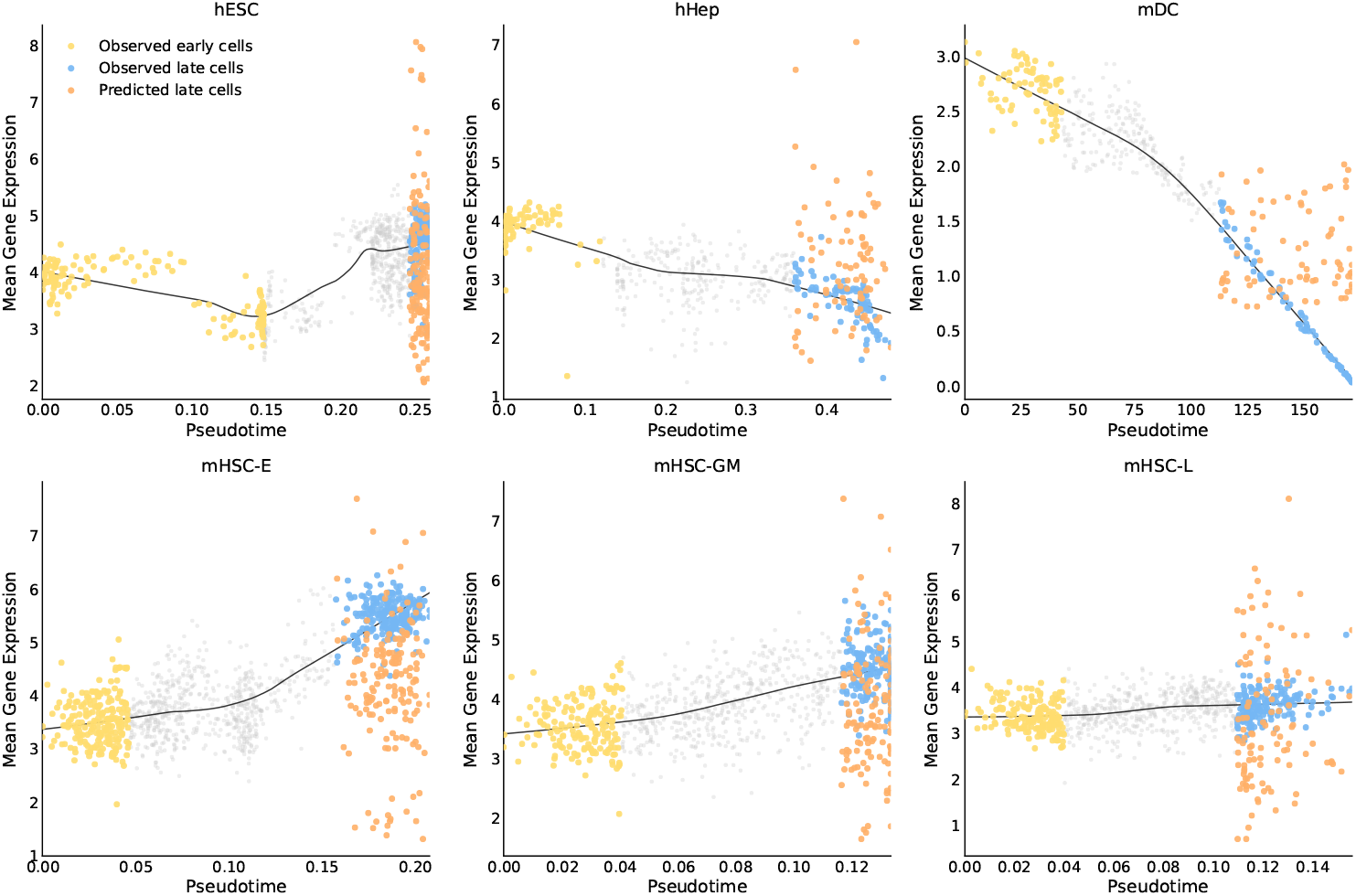
Dynamic transition probing across six datasets. Observed early cells, observed late cells, and predicted late states are shown along pseudotime, with the overall expression trend shown in gray.

We then compared dynamic probing performance across models and datasets. We excluded the mDC dataset from the aggregate comparison because its early-to-late expression changes were highly imbalanced, with nearly all evaluated genes being downregulated (Figure 8); this can bias directional accuracy toward trivial downregulation predictions. Across the remaining datasets, most single-cell foundation models achieved high directional accuracy (Figure 4(c)), suggesting that pretrained models contain useful signals for guiding expression changes. Notably, scFoundation showed the strongest overall performance, even though it was less competitive in static GRN reconstruction. This is consistent with a recent benchmark reporting that scFoundation can show relatively strong temporal separation and local velocity coherence in dynamical settings [40]. Together, these results suggest that static GRN recovery and dynamic transition probing capture different aspects of single-cell foundation models.

Next, we examined gene-level prediction behavior using scGPT on mHSC-L. Predicted expression changes were positively aligned with observed expression changes, indicating that the model captures the direction of many gene-level transitions rather than only shifting the global cell state (Figure 4(d)). Prediction accuracy was also higher for genes with larger observed changes (Figure 4(e)), suggesting that stronger dynamic signals are easier to recover, likely because they are less affected by noise and more constrained by learned gene-expression dependencies.

Finally, we tested whether this behavior depends on pretraining. We repeated the scGPT analysis with randomly initialized weights under the same iterative refinement procedure. Random weights produced much lower accuracy, whereas pretrained weights achieved substantially better performance (Figure 4(f)). This result indicates that the dynamic probing signal does not arise from the iterative procedure alone, but depends on regulatory and dynamical priors learned during pretraining.

## 4 Discussion

We introduced a unified benchmark for evaluating GRN reconstruction from single-cell foundation models, with dynamic transition probing as a complementary functional task. By comparing token embeddings, hidden states, and attention-derived scores under the same preprocessing, candidate gene sets, graph-construction rules, and reference networks, our framework separates representation quality from downstream implementation choices. The results reveal strong model-specific patterns: scGPT token embeddings provide strong zero-shot GRN signals, Geneformer benefits more from attention-derived scores, and scCello/LangCell often rely more on hidden-state representations. Thus, regulatory information is not stored in a universal model component, but depends on architecture and pretraining design.

Dynamic transition probing further shows that static GRN recovery and dynamic expression prediction capture different aspects of model knowledge. Several pretrained models can move early-cell states toward later states without temporal supervision, and scFoundation performs strongly in this dynamic setting despite weaker static GRN reconstruction. This divergence suggests that future benchmarks should evaluate both structural recovery and functional predictive capacity, rather than relying on static network reconstruction alone.

Our study has several limitations. First, reference GRNs are incomplete and biased toward well-studied genes; cell-type-specific ChIP-seq references yielded near-random performance for all methods, highlighting the limits of current ground truth. Second, we evaluate models in a zero-shot setting without fine-tuning, prompting, or task-specific adaptation, which may further improve regulatory recovery. Third, our extraction strategies are intentionally simple for fair comparison; more advanced approaches, such as head-selective attention aggregation or causal probing, may recover stronger signals. Finally, dynamic transition probing is currently focused on developmental trajectories, and extending it to perturbation response, disease progression, and multi-omic settings remains an important direction. Overall, our benchmark provides a controlled framework for studying how single-cell foundation models encode regulatory structure and expression dynamics, and may help guide the design of models that better connect static GRNs with dynamic cellular behavior.

## A Experimental Details

### A.1 Dataset processing

We followed the BEELINE preprocessing protocol for all six single-cell datasets used in the benchmark [7]. To select candidate genes for GRN inference, we quantified pseudotime-dependent expression variation using a generalized additive model for each gene and applied Bonferroni correction for multiple testing. Genes with corrected *P <* 0.01 were considered significantly variable. The numbers of selected genes, transcription factors (TFs), and network densities under different settings are summarized in Table 2.

**Table 2:**
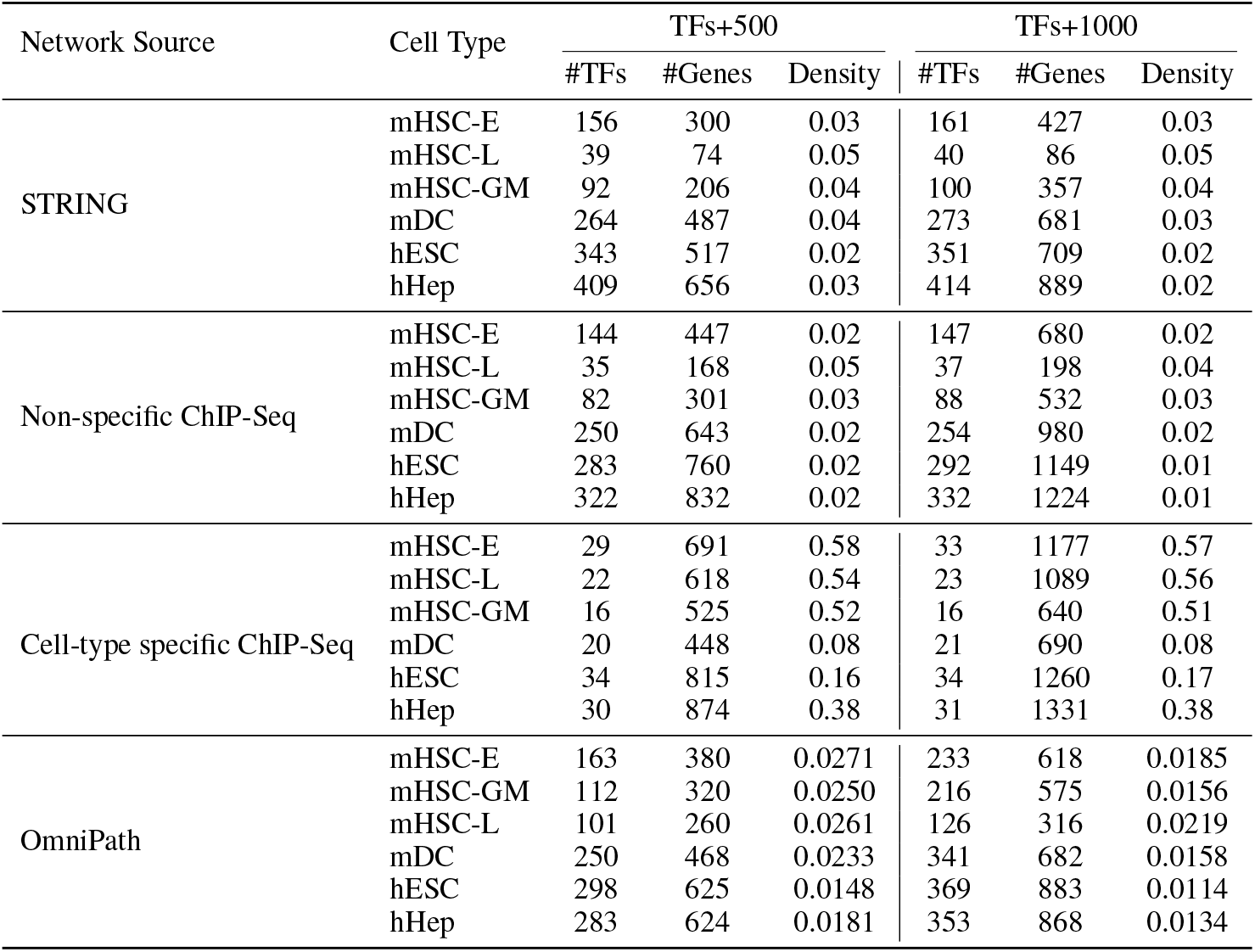
Statistics of regulatory networks under TFs+500 and TFs+1000 settings.

| Network Source | Cell Type | TFs+500 |  |  | TFs+1000 |  |  |
| --- | --- | --- | --- | --- | --- | --- | --- |
|  |  | #TFs | #Genes | Density | #TFs | #Genes | Density |
| STRING | mHSC-E | 156 | 300 | 0.03 | 161 | 427 | 0.03 |
|  | mHSC-L | 39 | 74 | 0.05 | 40 | 86 | 0.05 |
|  | mHSC-GM | 92 | 206 | 0.04 | 100 | 357 | 0.04 |
|  | mDC | 264 | 487 | 0.04 | 273 | 681 | 0.03 |
|  | hESC | 343 | 517 | 0.02 | 351 | 709 | 0.02 |
|  | hHep | 409 | 656 | 0.03 | 414 | 889 | 0.02 |
| Non-specific ChIP-Seq | mHSC-E | 144 | 447 | 0.02 | 147 | 680 | 0.02 |
|  | mHSC-L | 35 | 168 | 0.05 | 37 | 198 | 0.04 |
|  | mHSC-GM | 82 | 301 | 0.03 | 88 | 532 | 0.03 |
|  | mDC | 250 | 643 | 0.02 | 254 | 980 | 0.02 |
|  | hESC | 283 | 760 | 0.02 | 292 | 1149 | 0.01 |
|  | hHep | 322 | 832 | 0.02 | 332 | 1224 | 0.01 |
| Cell-type specific ChIP-Seq | mHSC-E | 29 | 691 | 0.58 | 33 | 1177 | 0.57 |
|  | mHSC-L | 22 | 618 | 0.54 | 23 | 1089 | 0.56 |
|  | mHSC-GM | 16 | 525 | 0.52 | 16 | 640 | 0.51 |
|  | mDC | 20 | 448 | 0.08 | 21 | 690 | 0.08 |
|  | hESC | 34 | 815 | 0.16 | 34 | 1260 | 0.17 |
|  | hHep | 30 | 874 | 0.38 | 31 | 1331 | 0.38 |
| OmniPath | mHSC-E | 163 | 380 | 0.0271 | 233 | 618 | 0.0185 |
|  | mHSC-GM | 112 | 320 | 0.0250 | 216 | 575 | 0.0156 |
|  | mHSC-L | 101 | 260 | 0.0261 | 126 | 316 | 0.0219 |
|  | mDC | 250 | 468 | 0.0233 | 341 | 682 | 0.0158 |
|  | hESC | 298 | 625 | 0.0148 | 369 | 883 | 0.0114 |
|  | hHep | 283 | 624 | 0.0181 | 353 | 868 | 0.0134 |

We considered two gene-selection strategies. First, we selected the top 500 or top 1000 dynamically variable genes among genes passing the corrected *P* -value threshold. Second, to avoid excluding TFs with modest expression variation but potential regulatory roles, we constructed two TF-prioritized gene sets. In the TFs+500 setting, all significant TFs were retained first, followed by the top 500 most dynamically variable non-TF genes. Similarly, TFs+1000 retained all significant TFs and added the top 1000 most dynamically variable non-TF genes. Unless otherwise stated, the main results are reported on TFs+500, with TFs+1000 results provided for comparison.

After applying each GRN inference method, we only evaluated edges whose source gene is a TF. This ensures that all predicted networks are compared under the same TF–target regulatory setting.

### A.2 Evaluated models and baselines

We evaluated six single-cell foundation models and three classical GRN inference baselines. The main architectural and pretraining characteristics of the foundation models are summarized in Table 3. Below, we briefly describe each model and baseline used in the benchmark.

**Table 3:** Architectural and pretraining characteristics of evaluated single-cell foundation models.

| Model | Attribute | Value |
| --- | --- | --- |
| scGPT | Model size | 50M |
|  | Pretraining cells | 33M |
|  | Gene embedding dim | 512 |
|  | Output dim | 512 |
|  | Vocabulary size | 60,697 |
|  | Layers | 12 |
|  | Heads | 8 |
|  | Architecture | Transformer encoder |
| Geneformer | Model size | 40M |
|  | Pretraining cells | 30M |
|  | Gene embedding dim | 512 |
|  | Output dim | 256 for 6-layer model; 512 for 12-layer model |
|  | Vocabulary size | 25,426 |
|  | Layers | 6 / 12 |
|  | Heads | 8 |
|  | Architecture | Transformer encoder with attention mask |
| scFoundation | Model size | 50M |
|  | Pretraining cells | 50M |
|  | Gene embedding dim | 768 |
|  | Output dim | 3072 |
|  | Vocabulary size | 19,264 |
|  | Layers | 40 |
|  | Heads | 16 |
|  | Architecture | Asymmetric encoder-decoder |
| LangCell | Model size | 40M |
|  | Pretraining cells | 40M |
|  | Gene embedding dim | 512 |
|  | Output dim | 256 |
|  | Vocabulary size | 25,426 |
|  | Layers | 12 |
|  | Heads | 8 |
|  | Architecture | Cell encoder initialized from Geneformer plus text encoder |
| scCello | Model size | 10M |
|  | Pretraining cells | 10M |
|  | Gene embedding dim | 256 |
|  | Output dim | 200 |
|  | Vocabulary size | 25,426 |
|  | Layers | 6 |
|  | Heads | 8 |
|  | Architecture | Cell-ontology-guided Transformer encoder |
| scPRINT | Model size | 50M |
|  | Pretraining cells | 45M |
|  | Gene embedding dim | 512 |
|  | Output dim | 512 |
|  | Vocabulary size | 30,000 |
|  | Layers | 12 |
|  | Heads | 8 |
|  | Architecture | Transformer encoder-decoder |

#### Geneformer

Geneformer [15] is a Transformer-based single-cell foundation model pretrained on the Genecorpus dataset. It represents each cell by ranking genes according to normalized expression and uses the top-ranked genes as input tokens. Geneformer provides 6-layer and 12-layer checkpoints; we used the larger checkpoint in our benchmark.

#### scGPT

scGPT [16] is a generative single-cell foundation model pretrained on large-scale single-cell omics data. It uses gene tokens together with cell-dependent binned expression values to encode input cells. We used the whole-human scGPT checkpoint pretrained on 33 million normal human cells.

#### ScFoundation

scFoundation [23] uses the asymmetric encoder–decoder architecture xTrimoGene to model 19,264 protein-coding genes. Unlike rank- or bin-based models, it projects continuous expression values into learnable value embeddings, preserving quantitative expression information. The model was pretrained on 50 million single cells using a read-depth-aware objective with source and target count tokens.

#### LangCell

LangCell [24] introduces a language–cell pretraining framework with a cell encoder and a text encoder initialized from Geneformer [15] and PubMedBERT [41], respectively. It uses Geneformer-like gene ranking and adds cross-modal cell–text contrastive learning objectives based on the scLibrary dataset.

#### ScCello

scCello [25] is a cell-ontology-guided single-cell foundation model pretrained on labeled scRNA-seq data mapped to a cell ontology graph. It uses rank-based gene expression encoding similar to Geneformer, while ontology-guided training encourages the model to capture hierarchical cell-type relationships. We used the default pretrained checkpoint released by the authors.

#### ScPRINT

scPRINT [26] is a Transformer-based foundation model for scRNA-seq data. Each gene representation combines gene identity, expression value, and genomic positional encodings. The model is pretrained on large-scale cellxgene data with objectives including denoising reconstruction, bottleneck reconstruction, and multi-label prediction. We used the released pretrained checkpoint for representation extraction and reconstruction-based dynamic probing.

#### GENIE3

GENIE3 [11] formulates GRN inference as a collection of tree-based regression problems, where the expression of each target gene is predicted from candidate regulators. Regulatory weights are computed from feature-importance scores. We used the standard GENIE3 implementation with default hyperparameters.

#### DeepSEM

DeepSEM [13] is a deep generative model for single-cell GRN inference. It combines a structural equation model with a regularized variational autoencoder to infer directed regulatory relationships from scRNA-seq data in an unsupervised manner.

#### GRNBoost

GRNBoost [12] is a scalable tree-based method for GRN inference from single-cell transcriptomic data. It uses gradient-boosted regression to predict target gene expression from candidate regulators and derives regulatory weights from feature importance. We used the default benchmark configuration.

## B Details of Static GRN Reconstruction

### B.2 Representation extraction strategies

#### Pretrained token embeddings

For each model, pretrained token embeddings are obtained from the gene embedding layer or vocabulary-aligned model parameters. Let **W**_emb_ ∈ℝ^*V* ×*d*^ denote the embedding matrix, where *V* is the vocabulary size and *d* is the embedding dimension. For gene token *i* with vocabulary index *x*_*i*_, its token embedding is

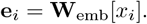

Because token embeddings are fixed for each gene and do not depend on a specific input cell, they represent static gene-level priors learned during pretraining. We compute pairwise cosine similarity between gene embeddings:

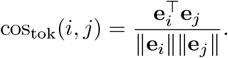

The resulting scores define an undirected gene–gene association network.

#### Final-layer hidden states

For hidden-state extraction, each input cell is passed through the model and the final-layer representation of each gene token is collected. Let **E** ∈ ℝ^*n*×*d*^ be the input embedding matrix for a cell and let

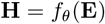

denote the final hidden-state matrix after the model forward pass. The hidden representation of gene *i* is **h**_*i*_ = **H**[*i*]. Unlike token embeddings, hidden states are input-dependent and encode contextual information from the full cell profile. We aggregate cell-wise hidden states into gene-level representations and compute pairwise cosine similarity:

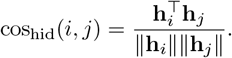

This produces an undirected GRN based on contextualized gene representations.

#### Attention-derived scores

For Transformer-based models, we also extract attention matrices as model-internal gene–gene coupling signals. For a given layer and head, the attention weight from gene *i* to gene *j* is computed from query and key vectors as

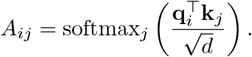

We aggregate attention weights of the last layer across heads and cells to obtain a single directed score for each ordered gene pair. Because attention is asymmetric, the score from gene *i* to gene *j* need not equal the score from gene *j* to gene *i*, yielding a directed GRN.

### B.2 GRN reconstruction

Each method outputs a ranked set of gene–gene scores in the form (Gene1, Gene2, weight). For fair comparison, all predictions are evaluated in a unified TF–target candidate space. The source gene must be a ground-truth TF, and the target gene must belong to the reference gene set. For undirected embedding-based networks, the gene–gene similarity score is assigned to the corresponding TF–target pair whenever one gene is a TF. For directed attention networks, the original source–target direction is retained.

Predicted edges are ranked by the absolute value of their edge weight, with ties assigned the same rank. For early-precision evaluation, we select top-ranked edges until the number of selected edges equals the number of edges in the corresponding ground-truth network. AUPRC and EPR are then computed within the same restricted candidate space.

### B.3 Evaluation methods

#### Edge-level evaluation

We evaluate edge recovery using Early Precision Ratio (EPR) and Area Under the Precision–Recall Curve ratio (AUPRC ratio). Candidate edges are restricted to TF–target pairs as described above. AUPRC measures the quality of the full ranked edge list and is computed using average precision. EPR focuses on the top-ranked predictions by comparing the precision among the top-*K* predicted edges, where *K* equals the number of ground-truth edges, against a random baseline. Formal definitions are provided in Appendix D.

#### TF-level evaluation

For each TF, we evaluate whether the predicted network recovers its target genes. Let *P* be the set of genes predicted as targets of a TF, and let *G* be the corresponding reference target set. Precision, recall, and F1 score are computed for each TF. We then rank TFs by F1 score and use the top 100 TFs as representative high-confidence regulators for overlap analysis and visualization. Consensus TFs are defined as TFs shared across the top-100 lists of all three scGPT extraction strategies, after excluding TFs with fewer than 20 predicted targets.

#### Topology-level evaluation

To evaluate topological fidelity, we compute five graph metrics: average shortest path length, clustering coefficient, modularity, average degree, and assortativity. Directed networks are first converted to undirected simple graphs. Specifically, if at least one directed edge exists between two genes in either direction, a single undirected edge is added; reciprocal edges are deduplicated, and self-loops are removed.

For each metric, we compare the predicted network with the STRING reference using a normalized closeness score:

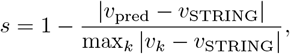

where *v*_pred_ is the metric value of a predicted network, *v*_STRING_ is the metric value of the STRING reference, and the denominator is the maximum deviation from STRING across all evaluated methods. This score maps the closest network to higher values, with STRING assigned a score of 1.

## C Details of Dynamic Transition Probing

### C.1 Continuous-value models

Continuous-value models, including scGPT, scFoundation, and scPRINT, operate directly on expression magnitudes and preserve quantitative differences across cells.

#### Problem setup

Let **x**^(*t*)^ ∈ℝ^*G*^ denote the expression state at iteration *t*, where *G* is the number of genes under evaluation. Early cells are defined by the lowest pseudotime quantile, and late cells by the highest pseudotime quantile. We use *q* = 20% in all experiments. The initial state **x**^(0)^ is the mean expression vector of early cells, and the observed early-to-late change is

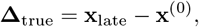

where **x**_late_ is the mean expression vector of late cells.

#### Iterative refinement

At each iteration, the current state is passed through a pretrained model to obtain a refined prediction:

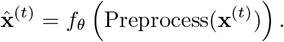

To avoid abrupt updates, we apply an exponential moving average:

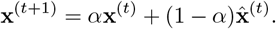

We set *α* = 0.9, corresponding to a 0.1 update rate for the model-predicted profile. Repeated application of this update generates a model-induced transition path

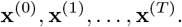

During inference, the model receives no time labels, pseudotime values, or late-cell profiles. The predicted expression change is

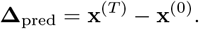

### C.2 Rank-driven models

Rank-driven models, including Geneformer, LangCell, and scCello, represent cells as gene-rank sequences rather than continuous expression vectors. This encoding preserves relative expression order but discards absolute expression magnitudes.

#### Problem setup

Let **r**^(*t*)^ ∈ℤ^*G*^ denote the rank-based token state at iteration *t*. The initial rank state **r**^(0)^ is obtained by ranking genes in each early cell from high to low expression and mapping the ranked genes to model-specific token IDs. Continuous expression values are used only for evaluation through **Δ**_true_.

#### Swap-based refinement

At iteration *t*, the current token sequence **r**^(*t*)^ is passed into the pretrained model to obtain token logits. We refine the sequence by proposing swaps between pairs of token positions while preserving the same set of expressed genes. For positions *i* and *j* with tokens 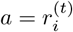 and 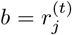, both positions are masked and the model compares the log-likelihood of the current assignment,

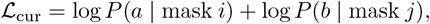

with the swapped assignment,

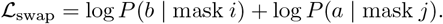

The swap is accepted if ℒ_swap_ *>* ℒ_cur_. We repeat this process for *T*_swap_ = 8 random swap proposals per iteration.

Because a lower rank position corresponds to higher expression, the predicted expression-direction signal from rank changes is

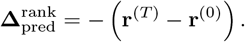

#### Directional accuracy

A gene is counted as directionally correct if the sign of the predicted change matches the sign of the observed early-to-late change. We report directional accuracy on the top 30% genes ranked by |**Δ**_true_|. Algorithm 1 summarizes the overall procedure.

##### Algorithm 1

Iterative refinement for dynamic transition probing

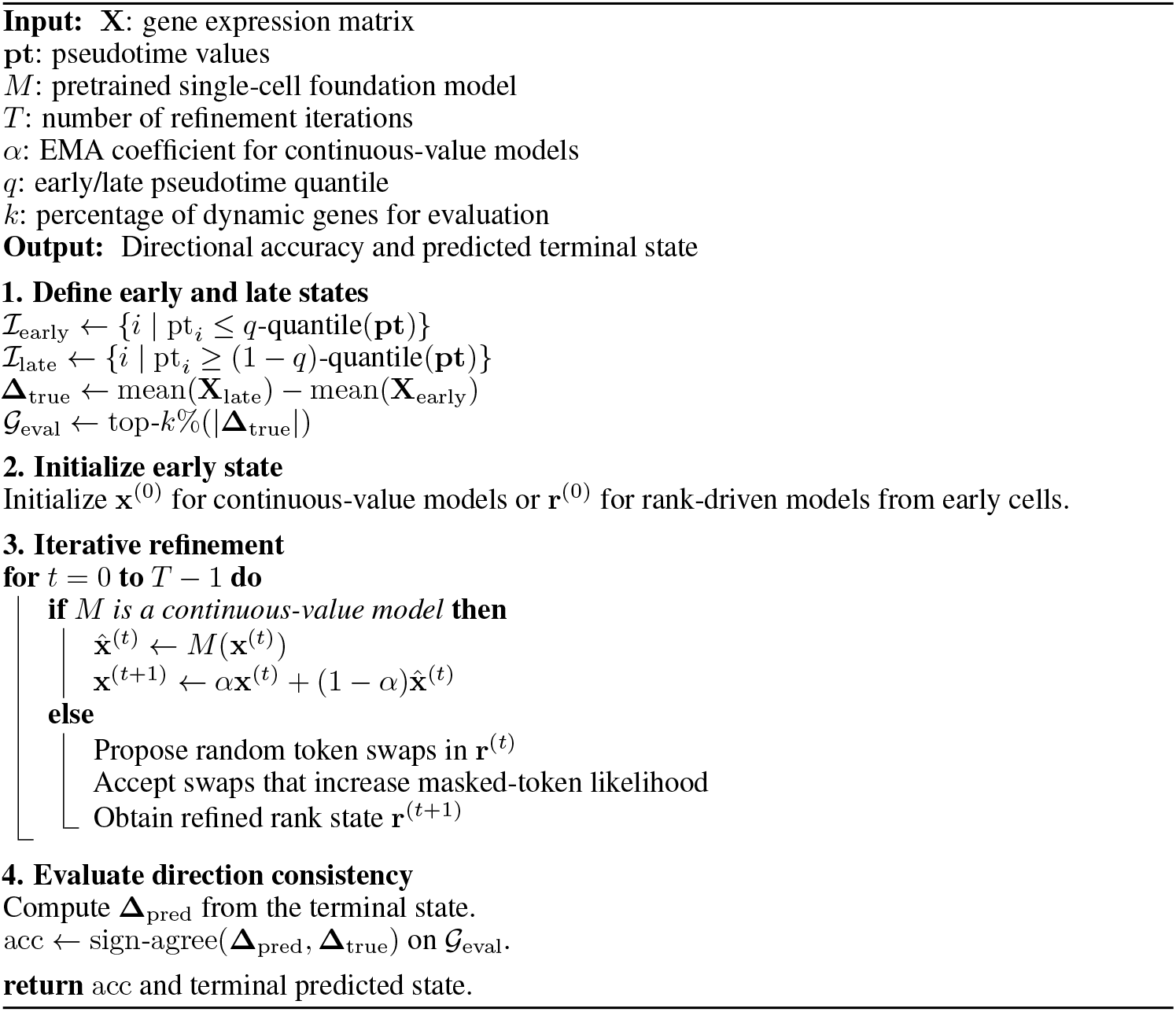

## D Evaluation Definitions

### D.1 Edge-level metrics

#### Early Precision Ratio (EPR)

Early precision measures the fraction of true positive edges among the top-*K* predicted edges, where *K* is set to the number of edges in the ground-truth network. Let EP denote this precision. The random baseline is the edge density within the restricted TF–target candidate space:

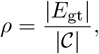

where *E*_gt_ is the set of ground-truth edges and is the set of all candidate TF–target pairs. The Early Precision Ratio is

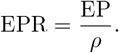

An EPR greater than 1 indicates enrichment over random ranking.

#### Area Under the Precision–Recall Curve ratio (AUPRC ratio)

For each threshold *θ*, precision and recall are defined as

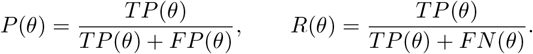

The AUPRC is the area under the precision–recall curve:

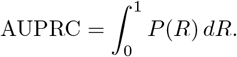

In the main results, we report the AUPRC ratio, defined as the observed AUPRC normalized by the random baseline precision *ρ*.

### D.2 TF-level F1 score

For each TF, we evaluate how well the predicted network recovers its target genes. Let *P* be the set of predicted targets for a TF and *G* be its reference target set. Precision and recall are

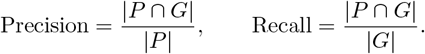

The F1 score is

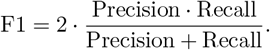

We compute F1 scores for all TFs and visualize their distributions using raincloud plots, where each point represents one TF. For each method, the top 100 TFs ranked by F1 score are used for overlap and downstream functional analyses.

### D.3 Topological metrics

All topology metrics are computed on undirected simple graphs. Directed networks are first projected to undirected graphs by ignoring edge direction, deduplicating reciprocal edges, and removing self-loops.

#### Average shortest path length

For networks with multiple connected components, we compute the average shortest path length on the largest connected component:

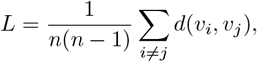

where *n* is the number of nodes in the largest connected component and *d*(*v*_*i*_, *v*_*j*_) is the shortest path distance between nodes *v*_*i*_ and *v*_*j*_.

#### Modularity

We apply Louvain community detection and compute modularity as

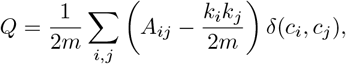

where *A*_*ij*_ is the adjacency matrix, *k*_*i*_ is the degree of node *i, m* is the number of edges, and *δ*(*c*_*i*_, *c*_*j*_) = 1 if nodes *i* and *j* belong to the same community.

#### Assortativity

Degree assortativity measures whether nodes preferentially connect to nodes with similar degree. We compute the degree assortativity coefficient using the NetworkX implementation. Positive values indicate assortative mixing, whereas negative values indicate disassortative mixing.

#### Average degree

Average degree measures overall network density:

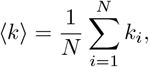

where *N* is the number of nodes and *k*_*i*_ is the degree of node *i*.

#### Clustering coefficient

For an undirected graph, the local clustering coefficient of node *i* is

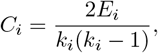

where *E*_*i*_ is the number of edges among the neighbors of node *i*. The average clustering coefficient is

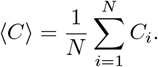

#### Degree distribution and CCDF

We characterize degree distributions using the complementary cumulative distribution function (CCDF):

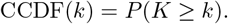

Empirically,

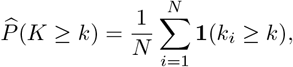

where *k*_*i*_ is the degree of node *i* and **1**(·) is the indicator function.

## E Complete Evaluation Results

Tables 4–7 report detailed per-dataset results for all methods across the six single-cell datasets. The aggregated mean ± standard deviation values in the main text are computed from these per-dataset results. Cell-type-specific ChIP-seq references are excluded from the main comparison because all methods showed near-random performance on this setting, but the complete results are still reported below for transparency.

**Table 4:**
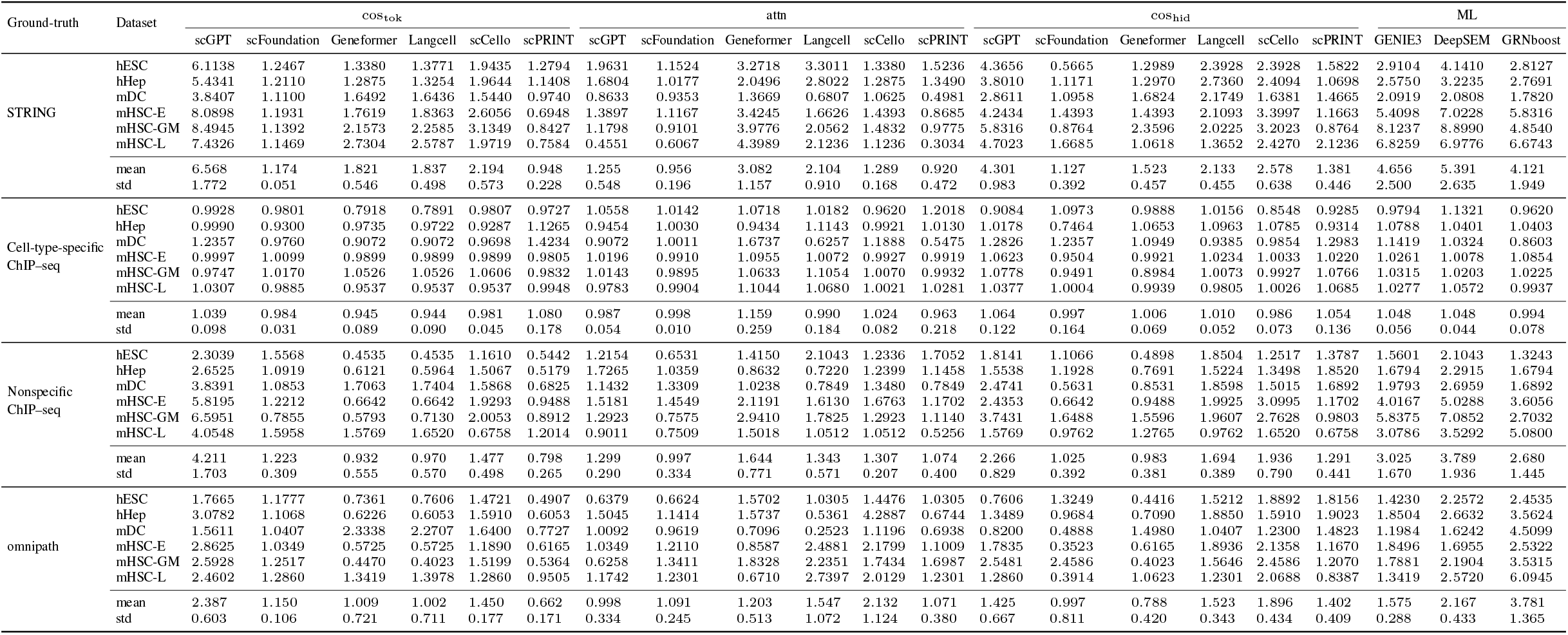
Early precision ratio of different methods (TF+500).

**Table 5:**
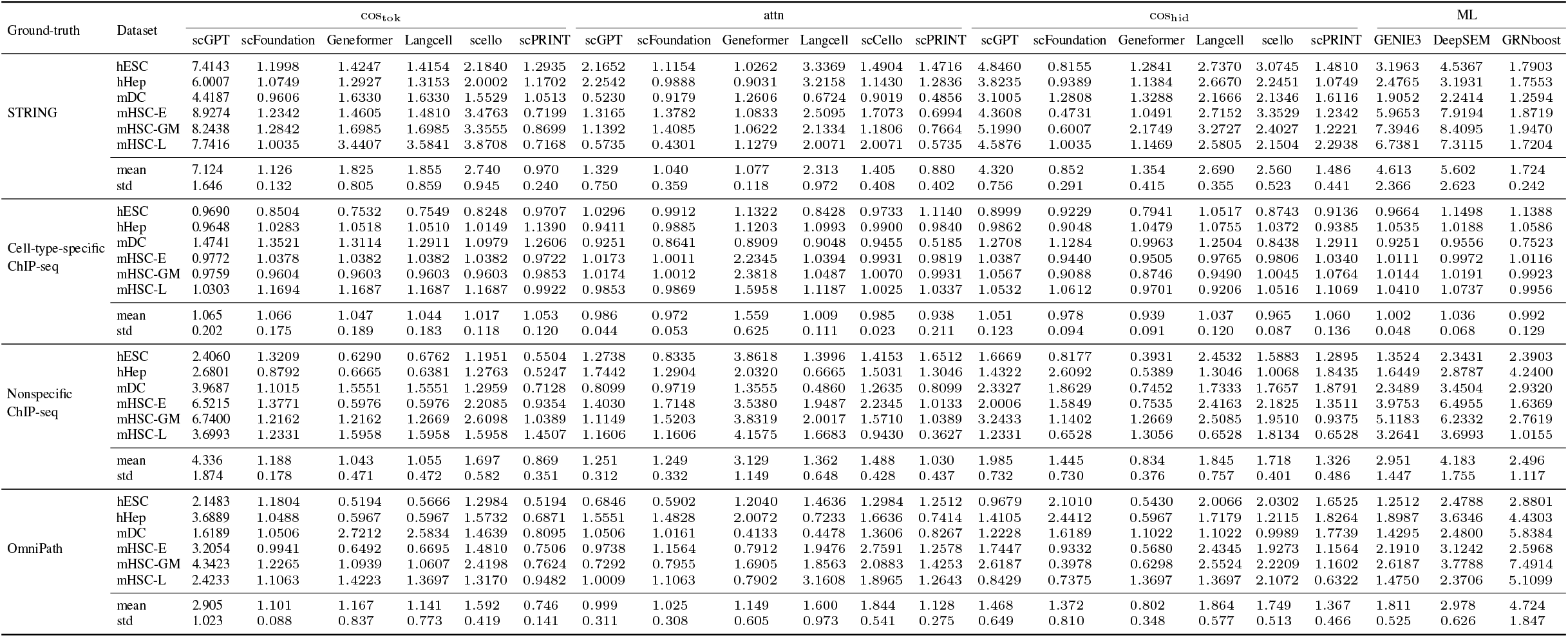
EPR Performance comparison of different methods across datasets (TF+1000).

**Table 6:**
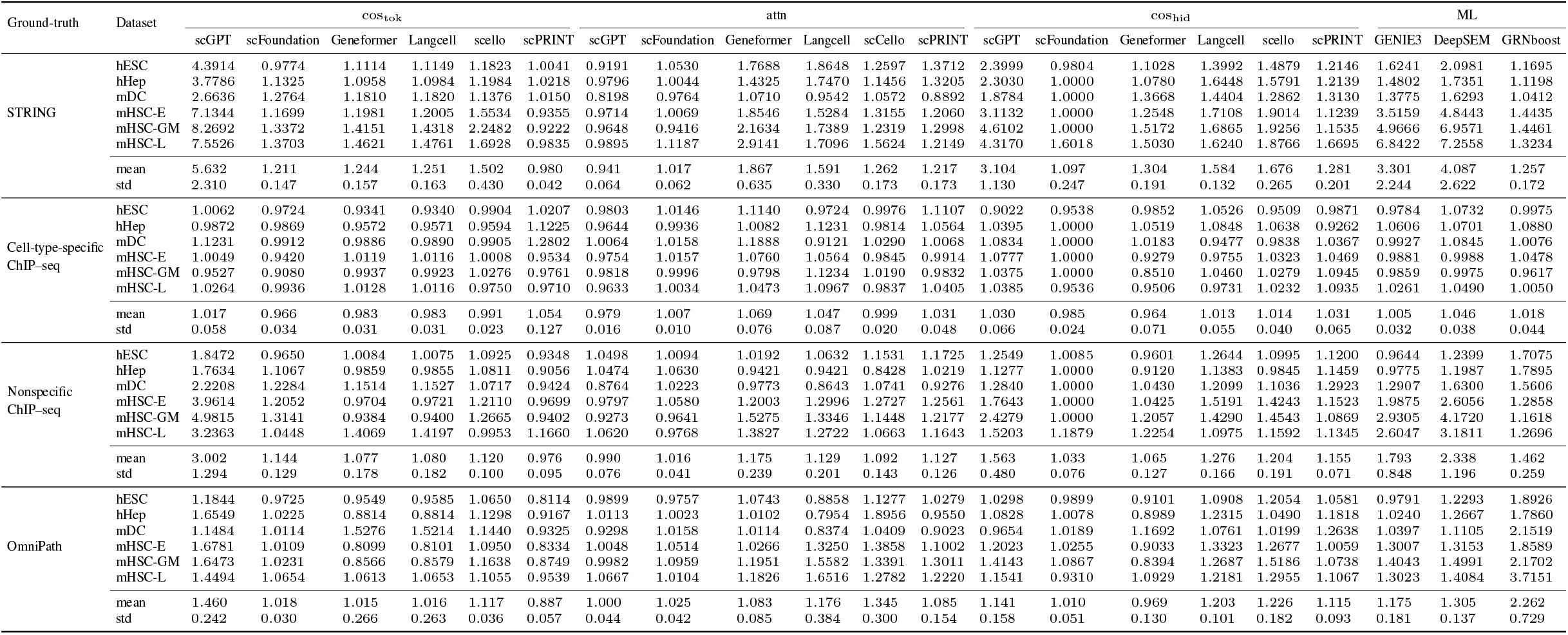
AUPR Performance comparison of different methods across datasets (TF+500).

**Table 7:**
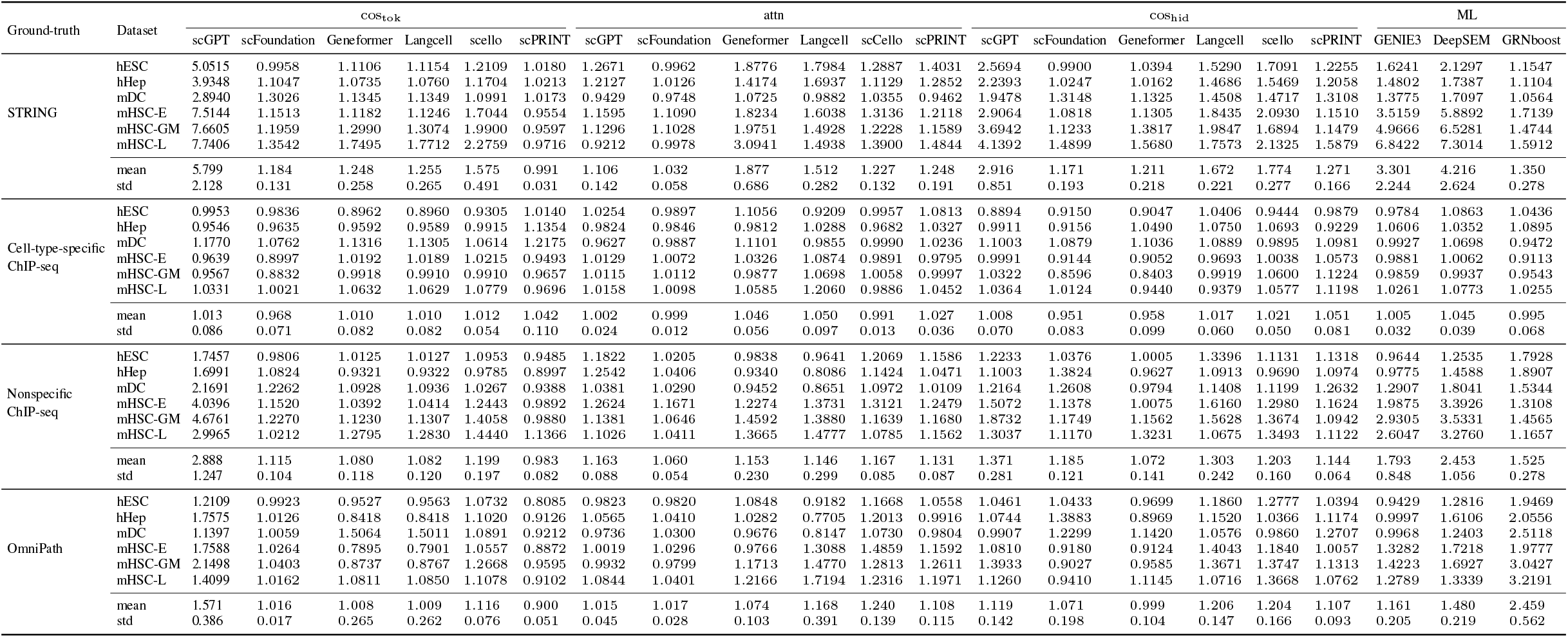
AUPRC Performance comparison of different methods across datasets (TF+1000).

**Table 8:**
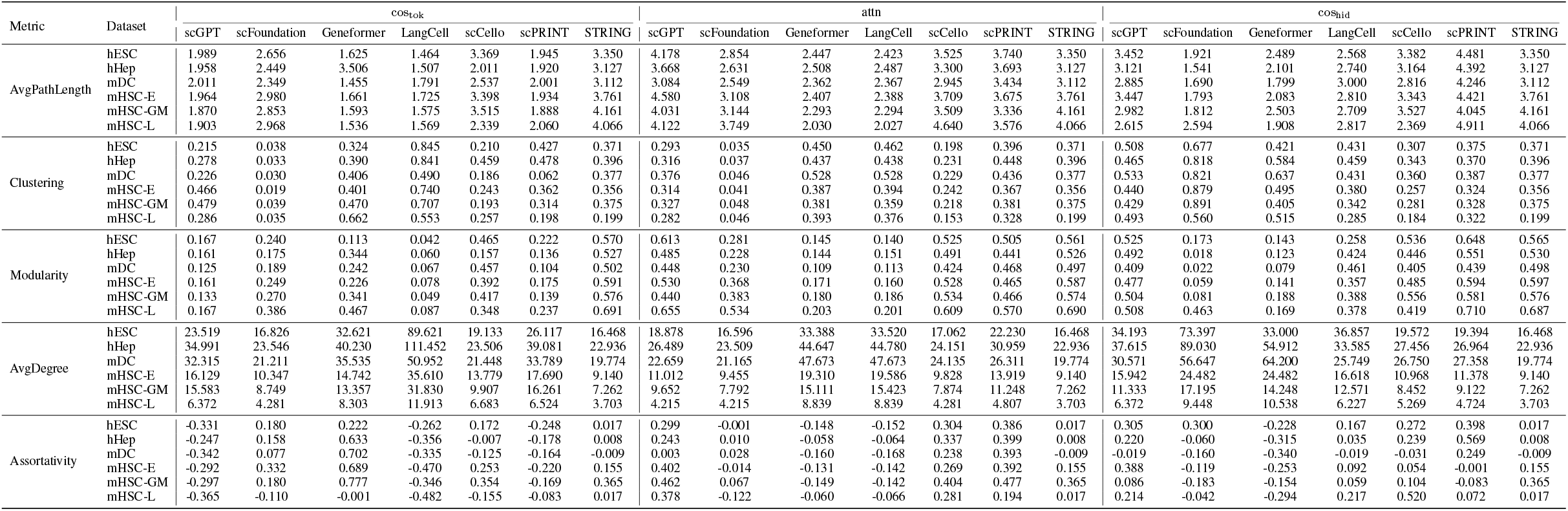
Network topology metrics for reconstructed GRNs across all datasets. Metrics are reported for networks reconstructed using cos_tok_, attn, and cos_hid_, together with the STRING reference network.

## F Further Experimental Results

To further examine the heterogeneity of attention heads, we analyzed topology metrics for networks reconstructed from individual scGPT attention heads on the hESC dataset. As shown in Figure 6, head-specific attention networks varied substantially in modularity, assortativity, and other topological properties. This suggests that different heads preserve different aspects of network structure, and that directly aggregating all heads may dilute useful topological signals.

**Figure 6:**
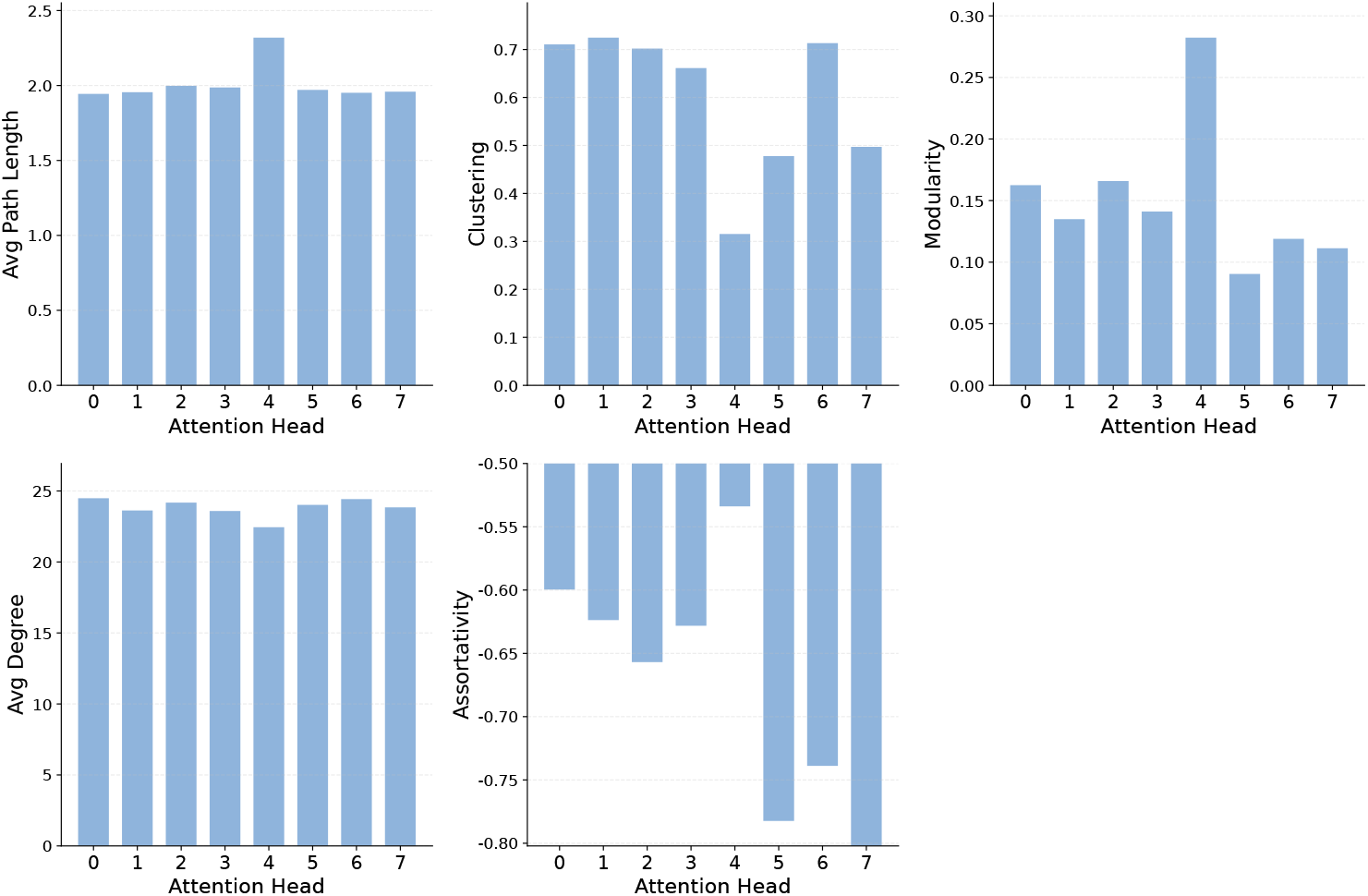
Topological metrics of scGPT GRNs reconstructed from individual attention heads on the hESC dataset. The results show substantial heterogeneity across heads.

We next examined whether the gene-level alignment between observed and predicted expression changes is consistent across datasets. As shown in Figure 7, predicted changes from scGPT were positively correlated with observed early-to-late expression changes in multiple datasets, although the strength of correlation varied by biological context. This supports the main-text conclusion that pretrained models can capture directional gene-level changes, but also shows that prediction quality is dataset dependent. The mDC dataset is a special case: because nearly all evaluated genes are downregulated from early to late states (Figure 8), its directional metrics may be biased and should be interpreted cautiously.

**Figure 7:**
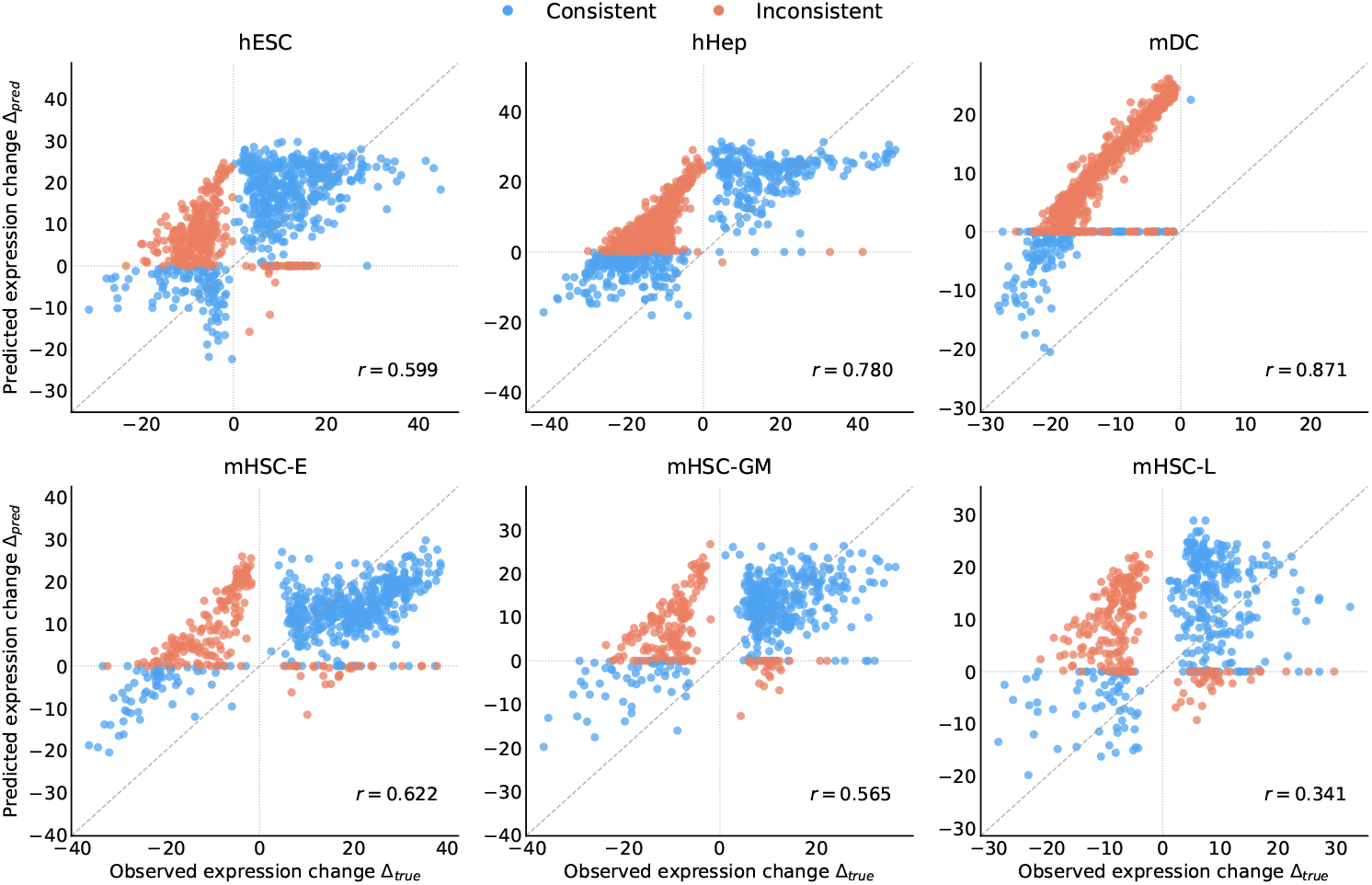
Relationship between observed expression changes and predicted expression changes for scGPT across all datasets. Each point represents one gene, and *r* denotes the Pearson correlation coefficient.

**Figure 8:**
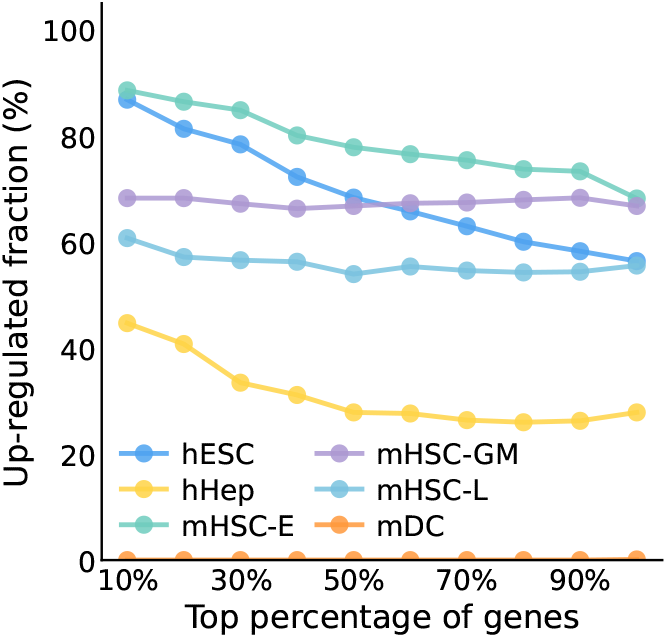
Percentage of genes with true up-regulation within progressively larger top-percentile subsets of genes, ranked by descending absolute observed expression change for six datasets.

Finally, we evaluated how directional accuracy changes over iterative refinement steps across all six datasets. As shown in Figure 9, most models showed stable or improved accuracy as refinement progressed, indicating that iterative updates generally improve dynamic transition probing. Performance varied across both models and datasets: scFoundation and Geneformer showed strong performance in several datasets, whereas scCello was less stable. These results support the main finding that dynamic transition probing depends on both model architecture and biological context.

**Figure 9:**
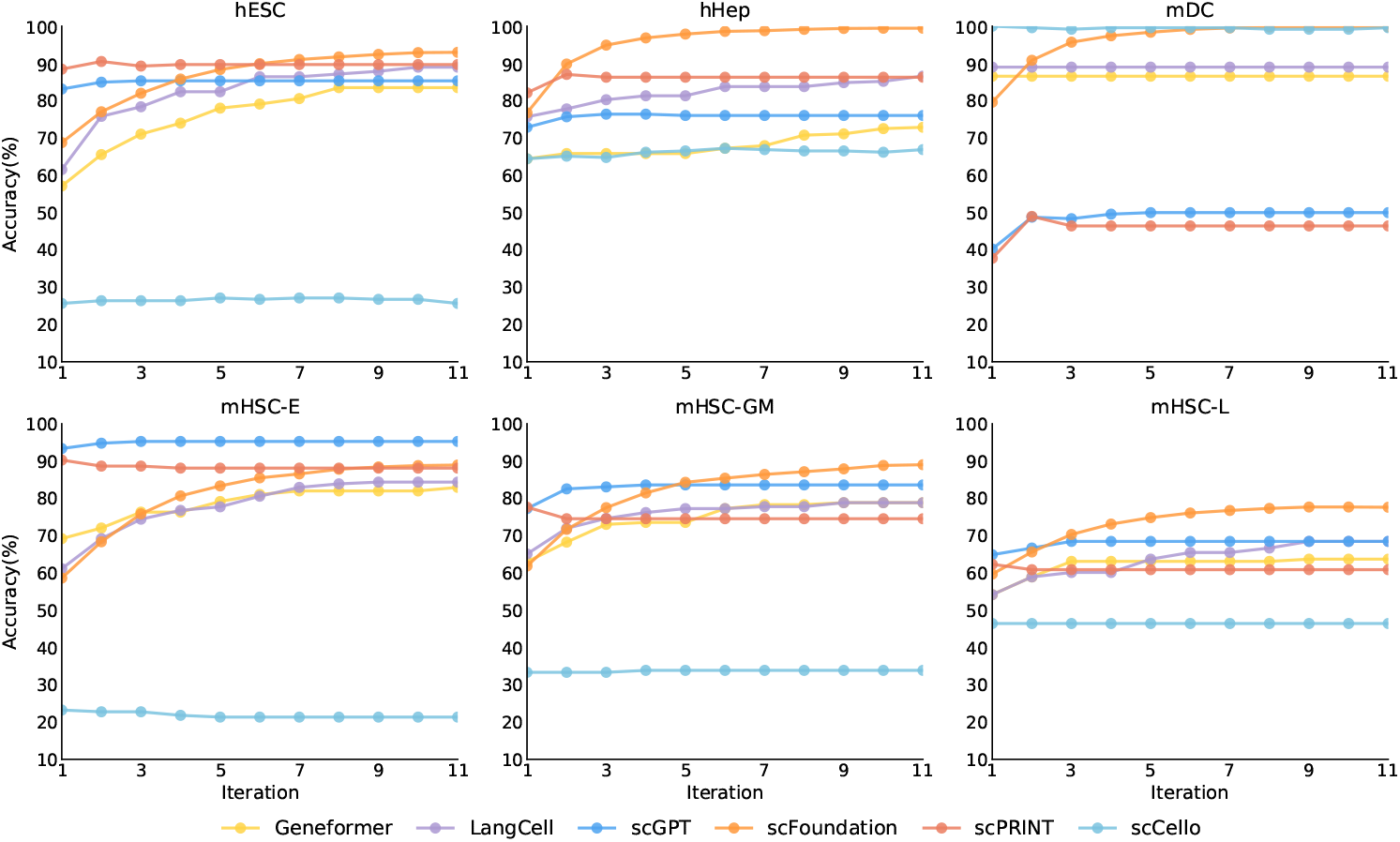
Directional prediction accuracy over iterative refinement steps across six datasets. Accuracy is computed on the top 30% genes with the largest observed early-to-late expression changes.

## Notes

### Competing Interest Statement

The authors have declared no competing interest.

## References

[1] Michael Levine and Eric H Davidson. Gene regulatory networks for development. Proceedings of the National Academy of Sciences, 102(14):4936–4942, 2005.

[2] Eric H Davidson and Michael S Levine. Properties of developmental gene regulatory networks. Proceedings of the National Academy of Sciences, 105(51):20063–20066, 2008.

[3] Mark W E J Fiers, Liesbeth Minnoye, Sara Aibar, Carmen Bravo González-Blas, Zeynep Kalender Atak, and Stein Aerts. Mapping gene regulatory networks from single-cell omics data. Briefings in Functional Genomics, 17(4):246–254, 2018.

[4] Pau Badia-i Mompel, Lorna Wessels, Sophia Müller-Dott, Rémi Trimbour, Ricardo O Ramirez Flores, Ricard Argelaguet, and Julio Saez-Rodriguez. Gene regulatory network inference in the era of single-cell multi-omics. Nature Reviews Genetics, 24(11):739–754, 2023.

[5] Richard Bonneau, David J Reiss, Paul Shannon, Marc T Facciotti, Leroy Hood, Nitin S Baliga, and Vesteinn Thorsson. The inferelator: an algorithm for learning parsimonious regulatory networks from systems-biology data sets de novo. Genome Biology, 7(5):R36, 2006.

[6] Hung Nguyen, Duc Tran, Bang Tran, Bahadir Pehlivan, and Tin Nguyen. A comprehensive survey of regulatory network inference methods using single cell RNA sequencing data. Briefings in Bioinformatics, 22(3):bbaa190, 2021.

[7] Aditya Pratapa, Ashwin P Jalihal, Jeremiah N Law, Adarsh Bharadwaj, and T M Murali. Benchmarking algorithms for gene regulatory network inference from single-cell transcriptomic data. Nature Methods, 17(2):147–154, 2020.

[8] Adam A Margolin, Ilya Nemenman, Katia Basso, Chris Wiggins, Gustavo Stolovitzky, Riccardo Dalla Favera, and Andrea Califano. Aracne: an algorithm for the reconstruction of gene regulatory networks in a mammalian cellular context. BMC Bioinformatics, 7(1):S7, 2006.

[9] Jeremiah J Faith, Boris Hayete, Joshua T Thaden, Ilaria Mogno, Jerome Wierzbowski, Guillaume Cottarel, Simon Kasif, James J Collins, and Timothy S Gardner. Large-scale mapping and validation of E. coli transcriptional regulation from a compendium of expression profiles. PLOS Biology, 5(1):e8, 2007.

[10] Anne-Claire Haury, Fabien Mordelet, Paola Vera-Licona, and Jean-Philippe Vert. Tigress: Trustful inference of gene regulation using stability selection. BMC Systems Biology, 6(1):145, 2012.

[11] Vân Anh Huynh-Thu, Alexandre Irrthum, Louis Wehenkel, and Pierre Geurts. Inferring regulatory networks from expression data using tree-based methods. PLOS ONE, 5(9):e12776, 2010.

[12] Thomas Moerman, Sara Aibar Santos, Carmen Bravo González-Blas, Jaak Simm, Yves Moreau, Stein Aerts, and Sara Aibar. Grnboost2 and arboreto: efficient and scalable inference of gene regulatory networks. Bioinformatics, 35(12):2159–2161, 2019.

[13] Hantao Shu, Jingtian Zhou, Qiuyu Lian, Han Li, Dan Zhao, Jianyang Zeng, and Jianzhu Ma. Modeling gene regulatory networks using neural network architectures. Nature Computational Science, 1:491–501, 2021.

[14] Jiaxing Chen, Chin Wang Cheong, Liang Lan, Xin Zhou, Jiming Liu, Aiping Lyu, William K Cheung, and Lu Zhang. Deepdrim: a deep neural network to reconstruct cell-type-specific gene regulatory network using single-cell rna-seq data. Briefings in Bioinformatics, 22(6):1–14, 2021.

[15] Constantine V Theodoris, Long Xiao, Aayush Chopra, Mark Chaffin, Zein Al Sayed, Mark Hill, Hannah Mantineo, Ethan M Brydon, Zexian Zeng, Wenjie Liu, et al. Transfer learning enables predictions in network biology. Nature, 618(7965):616–624, 2023.

[16] Haotian Cui, Jun Hong, Jiayuan Ding, Wenkai Zhang, Mingze Gao, et al. scgpt: Towards building a foundation model for single-cell multi-omics using generative ai. Nature Methods, 21:493–504, 2024.

[17] Daniel Marbach, James C Costello, Robert Küffner, Nicole M Vega, Robert J Prill, Diogo M Camacho, Kimberly R Allison, Manolis Kellis, James J Collins, and Gustavo Stolovitzky. Wisdom of crowds for robust gene network inference. Nature Methods, 9(8):796–804, 2012.

[18] Katarzyna Z Kędzierska, Lorin Crawford, Ava P Amini, and Alex X Lu. Zero-shot evaluation reveals limitations of single-cell foundation models. Genome Biology, 26(1):101, 2025.

[19] Takayuki Hayashi, Hiroki Ozaki, Yohei Sasagawa, Masahiro Umeda, Hiroki Danno, and Itoshi Nikaido. Single-cell full-length total rna sequencing uncovers dynamics of recursive splicing and enhancer rnas. Nature Communications, 9:619, 2018.

[20] Sonia Nestorowa, Fiona K Hamey, Blanca Pijuan Sala, Evangelia Diamanti, Mairi Shepherd, Elisa Laurenti, Nicola K Wilson, David G Kent, and Berthold Göttgens. A single-cell resolution map of mouse hematopoietic stem and progenitor cell differentiation. Blood, 128:e20–e31, 2016.

[21] J. Gray Camp, Keisuke Sekine, Tobias Gerber, Henry Loeffler-Wirth, Hans Binder, Martina Gac, Sabina Kanton, Joe Kageyama, Georg Damm, Daniel Seehofer, et al. Multilineage communication regulates human liver bud development from pluripotency. Nature, 546(7659):533–538, 2017.

[22] Li-Fang Chu, Ning Leng, Jue Zhang, Zhonggang Hou, Daniel Mamott, Wenjiang Ma, Tao Wang, Emily Zhong, Hongyu Jiang, Kenneth S Zaret, et al. Single-cell rna-seq reveals novel regulators of human embryonic stem cell differentiation to definitive endoderm. Genome Biology, 17:173, 2016.

[23] Minsheng Hao, Jing Gong, Xin Zeng, Chiming Liu, Yucheng Guo, Xingyi Cheng, Taifeng Wang, Jianzhu Ma, Xuegong Zhang, and Le Song. Large-scale foundation model on single-cell transcriptomics. Nature Methods, 21(8):1481–1491, 2024.

[24] Suyuan Zhao, Jiahuan Zhang, Yushuai Wu, Yizhen Luo, and Zaiqing Nie. Langcell: Language-cell pre-training for cell identity understanding. arXiv preprint 2405.06708, 2024.

[25] Xinyu Yuan, Zhihao Zhan, Zuobai Zhang, Manqi Zhou, et al. Cell ontology guided transcriptome foundation model. In Advances in Neural Information Processing Systems, pages 6323–6366. NeurIPS, 2024.

[26] J. Kalfon, J. Samaran, G. Peyré, et al. scprint: pre-training on 50 million cells allows robust gene network predictions. Nature Communications, 16:3607, 2025.

[27] S. Aibar, C.B. González-Blas, T. Moerman, et al. Scenic: single-cell regulatory network inference and clustering. Nature Methods, 14:1083–1086, 2017.

[28] Carrie A Davis, Benjamin C Hitz, Cricket A Sloan, Esther T Chan, Jean M Davidson, Idan Gabdank, Jason A Hilton, Kriti Jain, Ulugbek K Baymuradov, Aditi K Narayanan, et al. The encyclopedia of dna elements (encode): data portal update. Nucleic Acids Research, 46:D794– D801, 2018.

[29] Shinya Oki, Tazro Ohta, Go Shioi, Hisashi Hatanaka, Osamu Ogasawara, Yoshiki Okuda, Hideya Kawaji, Ryoko Nakaki, Jun Sese, and Chikara Meno. Chip-atlas: a data-mining suite powered by full integration of public chip-seq data. EMBO Rep., 19:e46255, 2018.

[30] Hua Xu, Caroline Baroukh, Ruth Dannenfelser, Emily Y Chen, Christopher M Tan, Yan Kou, Yong E Kim, Ihor R Lemischka, and Avi Ma’ayan. Escape: database for integrating high-content published data collected from human and mouse embryonic stem cells. Database, 2013:bat045, 2013.

[31] Luz Garcia-Alonso, Christian H Holland, Mahmoud M Ibrahim, Denes Turei, and Julio Saez-Rodriguez. Benchmark and integration of resources for the estimation of human transcription factor activities. Genome Research, 29:1363–1375, 2019.

[32] Zhi-Ping Liu, Canglin Wu, Hui Miao, and Hulin Wu. Regnetwork: an integrated database of transcriptional and post-transcriptional regulatory networks in human and mouse. Database, 2015:bav095, 2015.

[33] Heonjong Han, Hongseok Shim, Donghyun Shin, Jung Eun Shim, Yunhee Ko, Junha Shin, Hanhae Kim, Ara Cho, Eiru Kim, Tak Lee, et al. Trrust: a reference database of human transcriptional regulatory interactions. Scientific Reports, 5:11432, 2015.

[34] Damian Szklarczyk, Annika L Gable, David Lyon, Alexander Junge, Stefan Wyder, Jaime Huerta-Cepas, Milan Simonovic, Nadezhda T Doncheva, John H Morris, Peer Bork, et al. String v11: protein–protein association networks with increased coverage, supporting functional discovery in genome-wide experimental datasets. Nucleic Acids Research, 47:D607–D613, 2019.

[35] Dénes Türei, Tamás Korcsmáros, and Julio Saez-Rodriguez. Omnipath: guidelines and gateway for literature-curated signaling pathway resources. Nature Methods, 13(12):966–967, 2016.

[36] Michael P.H. Stumpf. Inferring better gene regulation networks from single-cell data. Current Opinion in Systems Biology, 27:100342, 2021.

[37] Weixu Wang, Zhiyuan Hu, Philipp Weiler, Sarah Mayes, Marius Lange, Jingye Wang, Zhengyuan Xue, Tatjana Sauka-Spengler, and Fabian J Theis. Regvelo: gene-regulatory-informed dynamics of single cells. bioRxiv, pages 2024–12, 2024.

[38] Jialong Jiang, Sisi Chen, Tiffany Tsou, Christopher S. McGinnis, Tahmineh Khazaei, Qin Zhu, Jong H. Park, Inna-Marie Strazhnik, Jost Vielmetter, Yingying Gong, John Hanna, Eric D. Chow, David A. Sivak, Zev J. Gartner, and Matt Thomson. D-spin constructs gene regulatory network models from multiplexed scrna-seq data revealing organizing principles of cellular perturbation response. bioRxiv, 2025.

[39] Albert-Laszlo Barabasi and Zoltan N Oltvai. Network biology: understanding the cell’s functional organization. Nature Reviews Genetics, 5(2):101–113, 2004.

[40] Xueya Zhou, Zihan Wang, Yue Ling, Qinxue Tian, Zhenyi Zhang, Yongge Li, Luonan Chen, and Peijie Zhou. Benchmarking zero-shot single-cell foundation model embeddings for cellular dynamics reconstruction, mar 2026.

[41] Qing Han, Shubo Tian, and Jinfeng Zhang. A pubmedbert-based classifier with data augmentation strategy for detecting medication mentions in tweets. arXiv preprint arXiv:2112.02998, 2021.

